# EGFR transactivates RON to drive oncogenic crosstalk

**DOI:** 10.1101/2020.08.11.246785

**Authors:** Carolina Franco Nitta, Ellen W. Green, Elton D. Jhamba, Justine M. Keth, Iraís Ortiz-Caraveo, Rachel M. Grattan, David J. Schodt, Aubrey C. Gibson, Ashwani Rajput, Keith A. Lidke, Mara P. Steinkamp, Bridget S. Wilson, Diane S. Lidke

## Abstract

**SUMMARY:** Crosstalk between disparate membrane receptors is thought to drive oncogenic signaling and allow for therapeutic resistance. EGFR and RON are members of two unique receptor tyrosine kinase (RTK) subfamilies that engage in crosstalk through unknown mechanisms. We combined high resolution imaging with biochemical studies and structural mutants to understand how EGFR and RON communicate. We found that EGF stimulation results in EGFR-dependent RON phosphorylation. Crosstalk is unidirectional, since MSP stimulation of RON does not trigger EGFR phosphorylation. Two-color single particle tracking captured the formation of complexes between RON and EGFR, supporting a role for direct interactions in propagating crosstalk. We further show that RON is a substrate for EGFR kinase, and transactivation of RON requires the formation of a signaling competent EGFR dimer. These results identify critical structural features of EGFR/RON crosstalk and provide new mechanistic insights into therapeutic resistance.

---

There is growing evidence demonstrating that crosstalk between members of distinct receptor tyrosine kinase (RTK) subfamilies can drive carcinogenesis and therapeutic resistance. Understanding these complicated interactions is critical for the development of novel dual-targeting therapeutics to improve patient outcomes^1–8^. Here we focus on the coordinated signaling between the Epidermal Growth Factor Receptor (EGFR, the canonical member of the EGFR/ErbB/Her subfamily) and Recepteur d’Origine Nantais (RON, also known as MSTR1 and a member of the Met subfamily). Prior evidence has implicated EGFR/RON crosstalk in the modulation of important cellular responses, notably migration and invasiveness in cancer^9–11^. RON expression combined with EGFR correlates with poorer outcomes for cancer patients. In head and neck cancer, EGFR/RON co-expression was associated with decreased event-free survival, while in bladder cancer, co-expression was correlated with increased tumor invasion, increased recurrence after first line therapy, and decreased patient survival^6^, ^9^. Direct interactions between RON and EGFR are supported by prior co-immunoprecipitation studies^6^, ^12^, as well as observations that EGFR/RON complexes can translocate into the nucleus to act as transcription factors^13^. These previous studies demonstrate EGFR/RON crosstalk, but lack significant details on the importance and complexity of this interaction.

Since the extracellular domains of EGFR and RON are so structurally distinct, it is difficult to explain their interactions through traditional dimerization models^14, 15^. EGFR is known to undergo receptor-mediated activation, where ligand binding induces a structural change exposing the dimerization arm to promote homo - and hetero-dimerization^16^. Like other members of the EGFR subfamily, EGFR relies on an asymmetric orientation of the kinase domains for activation^17^. Although the mechanisms of RON activation and potential dimerization are unknown, crystallographic studies of the RON extracellular domain have suggested that RON homodimers can form in the absence of ligand^14^.

Here, we combined high resolution imaging with rigorous biochemical measurements to dissect the mechanisms underlying EGFR/RON crosstalk and to contribute significantly to the understanding of their interactions. We provide evidence of unidirectional crosstalk between EGFR and RON. Activation of EGFR by EGF leads to RON phosphorylation via direct phosphorylation of RON by EGFR’s integral kinase which is further enhanced by RON’s own catalytic activity. Furthermore, EGFR activator or receiver mutants are incapable of promoting RON phosphorylation, demonstrating that RON cannot substitute for either partner of the EGFR asymmetric dimer. Taken together, our results support a molecular mechanism for crosstalk where EGF-bound EGFR forms heterotypic complexes with RON, independent of MSP, to promote RON activation and support RON-directed signaling outcomes.

## RESULTS

### Generation of human cell lines co-expressing full-length RON and EGFR

As model systems, we introduced RON into two well-characterized human cell lines, A431 and HEK-293. A431 cells have high levels of endogenous EGFR expression, providing a model system for tumors with high EGFR expression and modest levels of RON. HEK-293 cells, which have negligible levels of endogenous EGFR or RON, provided a test bed for balanced expression of combinations of RON plus either wildtype or mutant forms of EGFR. The low levels of endogenous RON expression in these cell lines allowed us to stably express full length HA-tagged RON (A431^RON^ and HEK^RON^), while avoiding potential complications from endogenous alternatively spliced RON isoforms^18–22^. ACP-tagged EGFR was also stably introduced into HEK^RON^ cells to generate a HEK-293 cell line expressing comparable levels of EGFR and RON (HEK^RON/EGFR^). Expression levels were evaluated by flow cytometry for both cell model systems. A431^RON^ cells display ~2.2 million EGFR molecules and only ~92,000 RON receptors on the cell surface (~24:1 EGFR:RON ratio), while HEK^RON/EGFR^ cells express EGFR and RON at a ratio of ~2:1 (~600,000 EGFR; ~275,000 RON). Confocal images show that RON and EGFR have a similar distribution on the plasma membrane of HEK^RON/EGFR^ (Figure 1A) and A431^RON^ cells (not shown).

**Figure 1.**
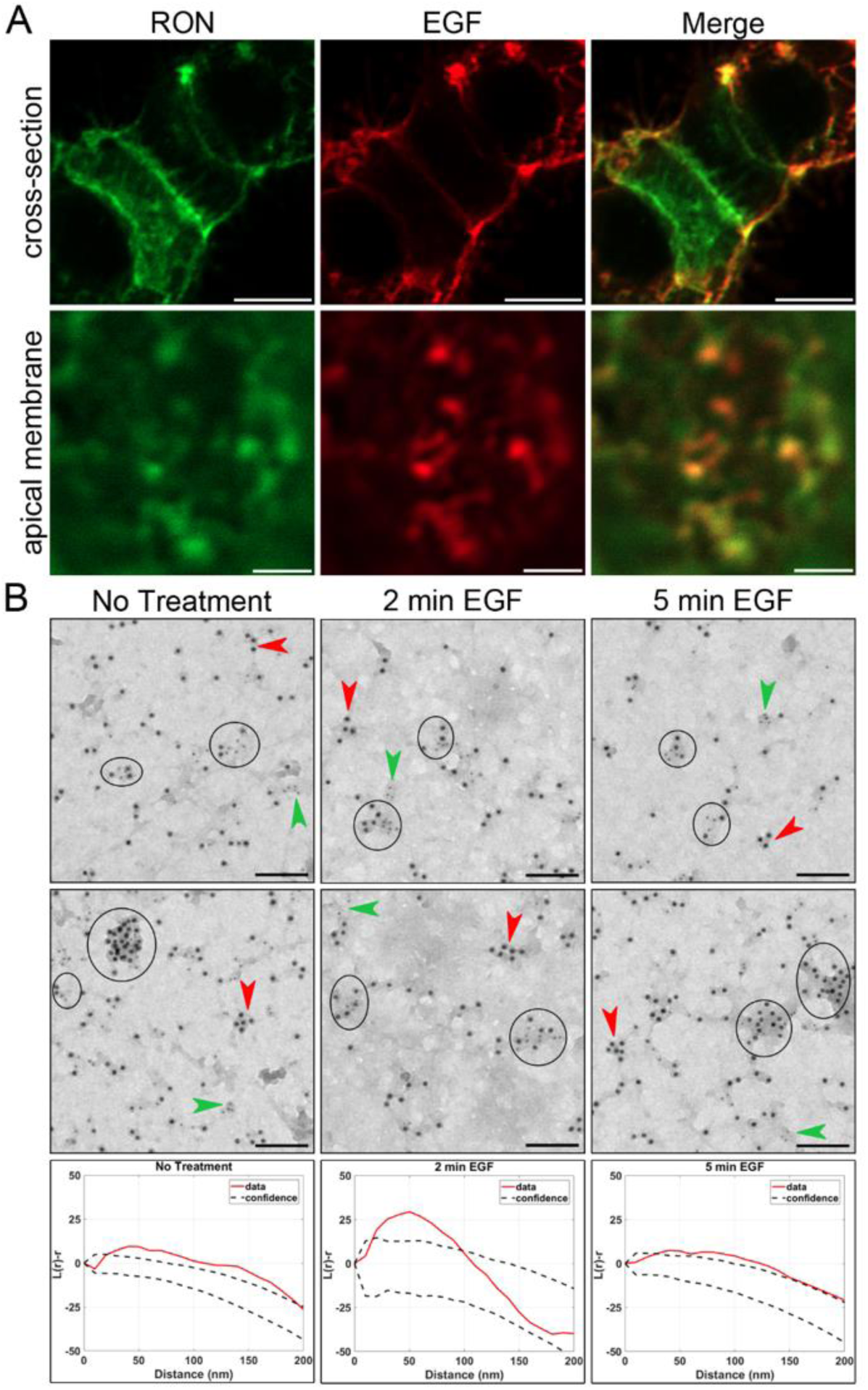
RON and EGFR co-culster in plasma membrane nanodomains. (A) HEK^RON/EGFR^ cells were first labeled for RON using α-HA-FITC Fab fragment(green), treated with 10 nM EGF-AF647 (red) for 5 min on ice and then fixed. Colocalization of RON and EGFR is seen at the plasma membrane. Scale bars, 10 μm (cross-section) and 2 μ (apical membrane). (B) Membrane sheets were prepared from A431^RON^ cells ± 50 nM EGF for 2 and 5 min. Sheets were labeled on the cytoplasmic face using amtibodies to RON (6 nm gold) and EGFR (12 nm gold). Circles indicate co-clusters of RON and EGFR; arrow heads indicate clusters containing RON (green) or EGFR (red) only. Bottom row: Ripley's K co-variant statistical test corresponding to the EM image immediately above. Scale bar, 100nm.

### RON and EGFR co-cluster in plasma membrane nanodomains

We applied our established transmission electron microcopy (TEM) technique with immunogold-labeled membrane sheets^23^ to evaluate the nano-organization of RON and EGFR. Receptor spatial distributions were determined from resting or EGF-stimulated A431^RON^ cells and imaged by TEM (Figure 1B). TEM images show that RON and EGFR frequently co-reside in mixed clusters in untreated cells (circles, Figure 1B, **left panels**). The close proximity of the two receptors on resting membranes was confirmed by Ripley’s K co-variant statistical test^23, 24^ (Figure 1B, **bottom panels**). EGFR/RON co-clustering is maintained after 2 and 5 min of treatment with 50 nM EGF (Figure 1B, **middle and right panels**).

### Crosstalk between EGFR and RON is EGF-driven

Given their strong co-localization, we evaluated EGFR/RON crosstalk based on changes in receptor phosphorylation in response to each of their cognate ligands. EGF treatment led to the expected EGFR phosphorylation in both A431^RON^ and HEK^RON/EGFR^ cells (Figure 2A), while MSP treatment induced RON phosphorylation (Figure 2B). Note that our western blot analysis resolved the mature RON (bottom RON band) from the pro-form (upper band; see **Figure S1A**). Importantly, treatment of cells with EGF promoted phosphorylation of RON (Figure 2B). This effect was dose-dependent and detectible at doses of EGF as low as 2 nM (**Figure S1B**). In contrast, neither physiological levels nor high doses of MSP could induce EGFR phosphorylation at PY1068 or other EGFR phospho-tyrosine sites (Supplemental **Figures S1C-D**). This was the first indication that crosstalk is unidirectional in our two model systems, with crosstalk occurring from EGF-bound EGFR to RON but not from MSP-bound RON to EGFR.

**Figure 2.**
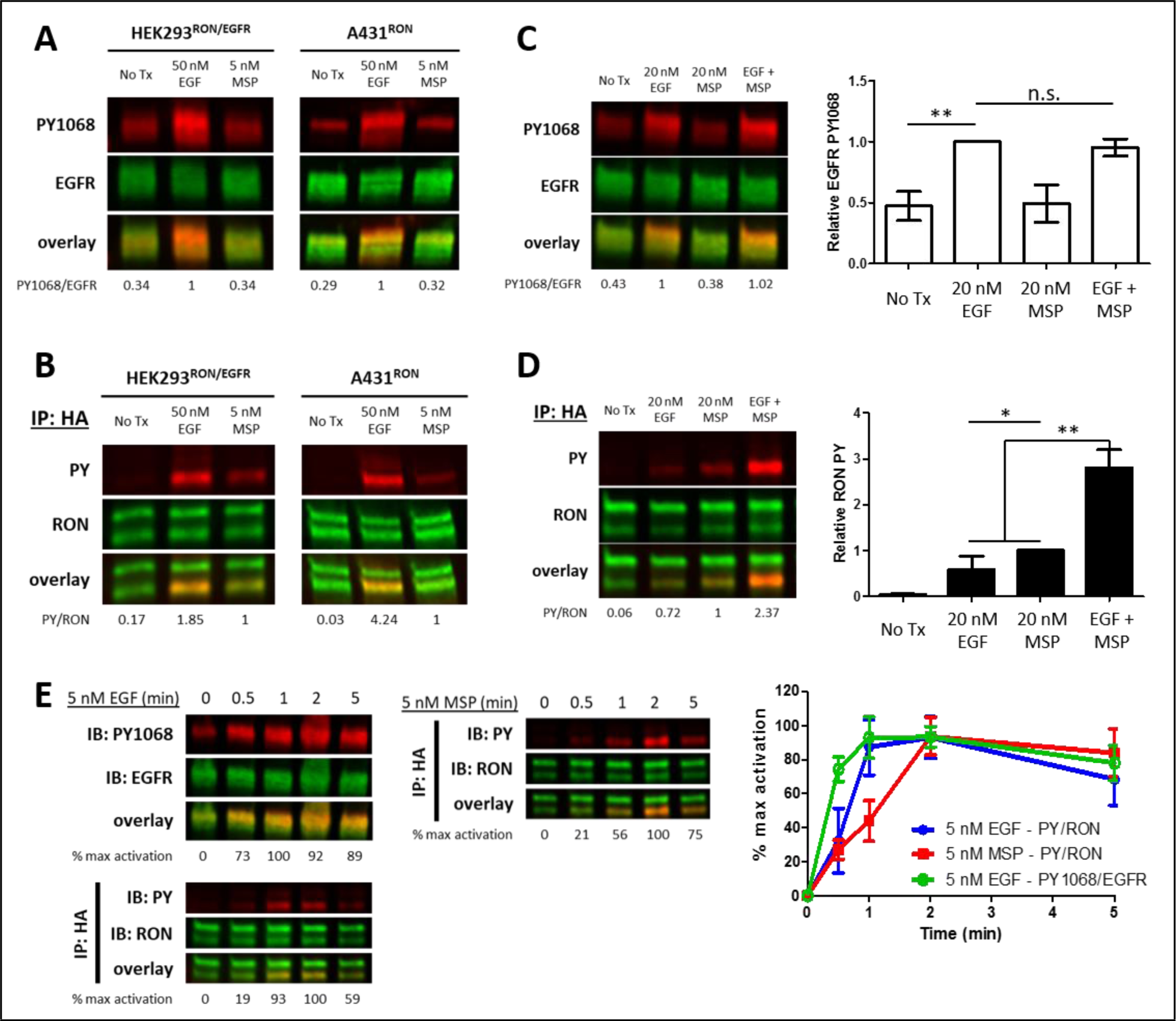
Crosstalk between EGFR and RON is EGF-driven. (A-B) HEK^RON/EGFR^ or A431^RON^ cells were treated with ±5 nM MSP or 50 nM EGF for 5 min at 37°C. Representative immunoblots showing PY 1068 and EGFR on cell lysates (A) or pan pan-phosphotyrosine (PY) and RON on samples immunoprecipitated (IP) with anti-HA antibody (B). (C-D) A431^RON^ cells were stimulated with ± 20 nM EGF, 20 nM MSP or both for 5 min at 37 °C and immunoblotted as in A and B. Triplicate experiments quantified in the bar graphs to the right, shown as mean ± SD. (E) Representative immunoblots of a phosphorylation time course for A431^RON^ cells treated with 5nM EGF or 5nm MSP and immunoblotted as in A and B. Graphed values (right) are from triplicate experiments, normalized to maximal activation, and presented as mean ± SD. * p <0.05; **p<0.01.

Dual stimulation with EGF and MSP did not increase EGFR phosphorylation (Figure 2C). However, the combination of EGF and MSP led to a synergistic enhancement in RON phosphorylation that is higher than would be expected from the additive effects of either ligand alone (Figure 2D). These results further support the conclusion that, while crosstalk occurs between full-length RON and EGFR, it is unidirectional and EGF-driven. EGFR was often detected in RON immunoprecipitates as a band co-migrating with pro-RON at 180 kDa (using EGFR or EGFR-PY1068 antibodies). However, co-precipitation is ligand-independent, as we show here and as previously reported^6, 12^ (**Figure S1A** and **Table S1**). Taken together with the observations in Figure 1 that co-localization is also ligand-independent, these data suggest that pre-existing protein complexes may contribute to EGFR-to-RON crosstalk.

### RON phosphorylation kinetics are accelerated with EGF stimulation

We next evaluated the phosphorylation kinetics of RON and EGFR in response to either MSP or EGF at similar doses. EGF-induced EGFR-PY1068 phosphorylation was rapid, peaking between ^30–60^ seconds (Figure 2E; **top left blot and green line**), as previously demonstrated^25–27^. RON phosphorylation after EGF treatment was similarly rapid, reaching maximum phosphorylation levels by 60 sec (Figure 2E; **bottom left blot and blue line**). In contrast, RON phosphorylation in response to its cognate ligand, MSP, was markedly slower, peaking at 2 min (Figure 2E; **right blot and red line**). The faster kinetics of EGF-driven RON phosphorylation, when compared to MSP-driven RON phosphorylation, led us to postulate that RON is a substrate for EGFR kinase activity in response to EGF binding.

### Crosstalk occurs at the plasma membrane

The evidence for co-localization at the plasma membrane and rapid unidirectional crosstalk, suggested that RON and EGFR form hetero-oligomeric complexes to alter EGF-driven signaling output. Using single particle tracking (SPT) of Quantum Dot (QD) labeled receptors we evaluated the mobility of HA-RON on the surface of live A431^RON^ cells using a monovalent anti-HA Fab fragment conjugated to QD probes (QD605-HA-RON)28. Our previous work has shown that mobility is a read-out for receptor phosphorylation states, such that a shift to slower mobility is correlated with receptor phosphorylation and recruitment of downstream signaling molecules^29^. Figure 3A shows the mean squared displacement (MSD) versus time lag (Δt) for tracking of QD605-HA-RON under different stimulation conditions. The average diffusion coefficients are given in Figure 3B. Consistent with ligand-induced phosphorylation and/or oligomerization, we observed that RON mobility is decreased upon MSP stimulation (Figure 3A-B, **blue**). Notably, RON mobility is also decreased with EGF addition (Figure 3A-B, **green**). This EGF-induced mobility change was prevented when cells were treated with EGFR kinase inhibitor (PD153035) (Figure 3A-B, **red**). Confocal images indicate the location of RON (green) and EGF (red) in HEK^RON/EGFR^ cells after 10 min of EGF stimulus (Figure 3C). As expected, EGF-bound EGFR is rapidly endocytosed and shows obvious co-localization with the early endosome marker, EEA1 (blue). In contrast, RON receptors are retained at the plasma membrane. The lack of co-endocytosis is further supported by TEM images from A431^RON^ cells, where EGFR, but not RON, was found in clathrin-coated pits 5 min after EGF addition (Figure 3D). These data support the premise that EGFR-mediated activation of RON occurs at the plasma membrane, rather than from endosomes, and is dependent on EGFR kinase activity. In addition, they suggest that EGFR/RON interactions are sufficiently transient that EGFR is sorted for endocytosis, while RON remains on the surface.

**Figure 3.**
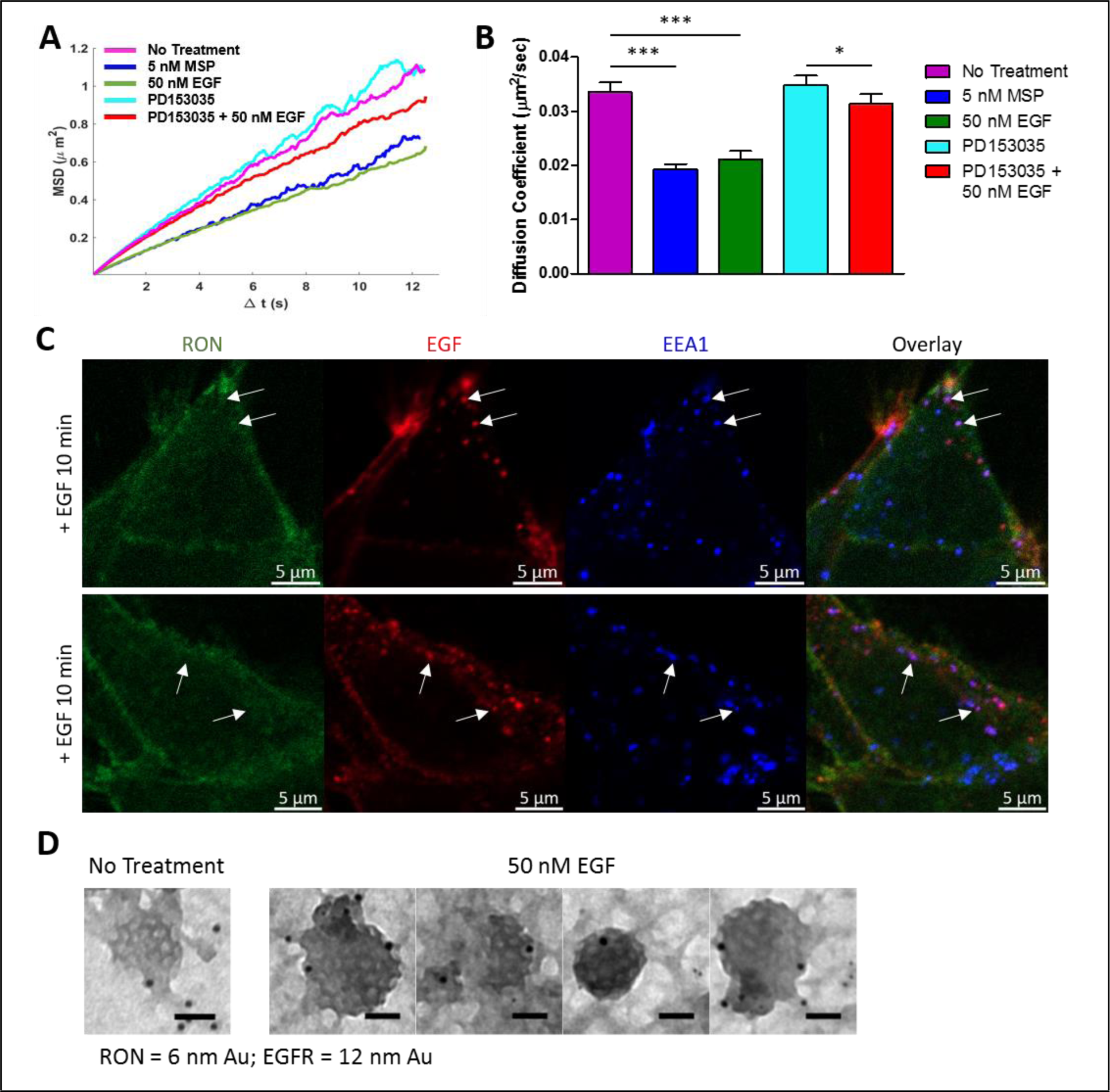
Crosstalk occurs at the plasma membrane. (A) Single particle tracking of QD605-HA-RON was used to quantify RON mobility on A431^RON^ cells ± Ensemble mean squared displacement (MSD) is plotted for n>1317 ttajectories per condition. A reduction in slope of the MSD indicates a reduced mobility. (B) Corresponding diffusion coeffcients for A given as mean ± SD. ^&^ p < 0.05; ^&&&^ p μ 0.001. (C) HEK^RON/EGFR^ cells were labeled for RON with α-HA-FITIC Fab fragment (green), treated with 10 nM EGF-AF647 (red) for 5 min on ice followed by 10 min at 37 °C, then fixed and labeled with an antibody to EEA1 (early endosomes, blue). EGF-positive endosomes (arrows) do not contain RON. (D) Membrane sheets prepared from A431^RON^ cells ± 50 nM min were labeled for RON (6 nm gold) or EGFR (12 nm). TEM images show clathrin-coated pit lattices on the cell membranes containing EGFR, but not RON. Scale bars, 50nm.

### EGF-bound EGFR and RON engage in direct interactions

To confirm that EGFR and RON interact at the cell membrane, we used simultaneous two-color QD tracking that allows for the direct detection and quantification of protein-protein interactions on live cells, as we have described previously^28–31^. Figure 4 demonstrates the visualization of receptor interactions by tracking of individual receptors in spectrally distinct channels at high spatiotemporal resolution. QDs were conjugated to either a monovalent anti-HA Fab fragment^28, 30^ for RON (QD-HA-RON) or EGF^29, 32^ to follow ligand-bound EGFR (QD-EGF-EGFR). We monitored RON/RON homo-interactions in A431^RON^ cells by labeling receptors with a mixture of anti-HA Fab conjugated to either QD605 or QD655 (Figure 4A). Figure 4B and **Video 1** shows an example of a long-lived interaction between two QD-tagged RON receptors that lasts for ~ 5 sec before breaking apart. A range of dimer lifetimes was observed, and additional examples of RON homo-interactions are found in **Figure S2A,B** and **Videos 2-3**. Two-color tracking was next used to determine if RON and EGFR form hetero-oligomeric complexes. Here, RON was tracked using anti-HA Fab conjugated to QD655, while endogenous EGFR is tracked with QD605-EGF (Figure 4D). This live cell imaging approach directly captures pairs of QD-labeled RON and EGF-bound EGFR that engage as complexes and move with correlated motion on the cell membrane. The example in Figure 4E and **Video 4** shows a more transient interaction with a duration of ~ 1.5 sec (see further examples in **Figure S2C,D and Videos 5-6**).

**Figure 4.**
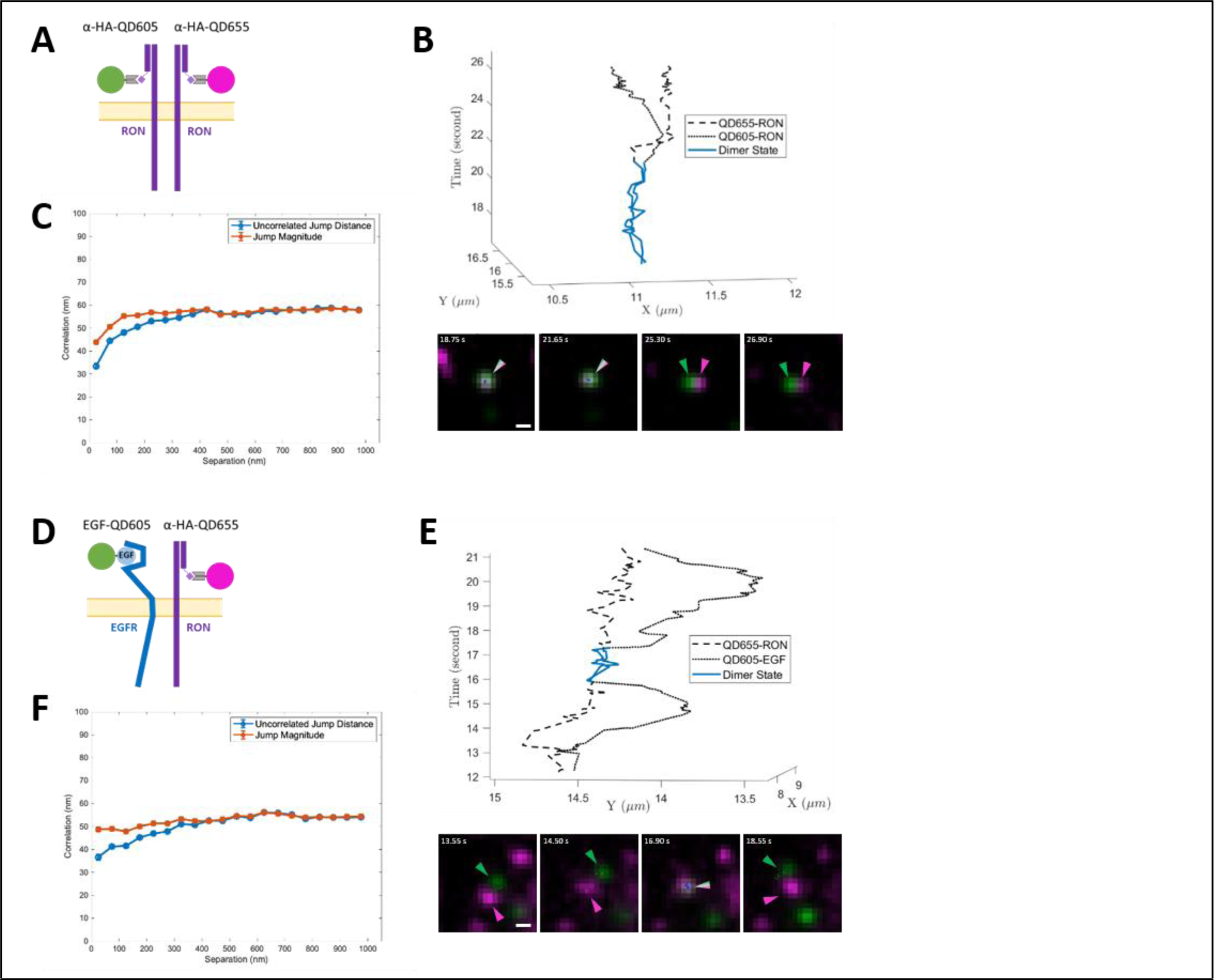
Two-color single QD tracking captures interactions between RON and EGFR. Two color SPT results for resting RON receptor interactions (A-C) and ligand-bound EGFR interaction with RON (D-F). (A) Schematic representation of two-color (QD605-HA-Fab and QD655-HA-Fab) RON SPT. (B) Representative time series (bottom) and 3D trajectory (top) for RON homo-interaction lasting ~5 sec (blue). Scale bar, 500 nm. (C) Ensemble correlated motion plot for all two-color RON tracking. (D) Schematic representation of two-color SPT of EGF-bound EGFR (QD655-EGF) and RON (QD605-HA-Fab). (E) Sample time series (bottom) and 3D trajectory(top) showing EGF-EGFR and RON interacting for ~1.5 sec (blue). Scale bar, 500 nm. (F) Ensemble corrrelated motion plot for all EGF-EGFR and RON tracking.

Quantification of correlated motion between receptors confirmed the formation of *bona fide* receptor complexes^29^. The presence of correlated motion is assessed over the full data-set of two-color trajectories and, therefore, reports on the behavior of the overall population. Correlated motion was observed when two RON receptors are in close proximity, as indicated by the reduction in the uncorrelated jump distance at small separation seen in Figure 4C. Jump magnitude also decreases at small separation, indicating that RON homo-complexes are moving more slowly than monomers. Importantly, correlated motion is clearly observed for RON and EGF-bound EGFR, confirming direct interactions between these disparate receptors (Figure 4F).

Using a two-state hidden Markov model (HMM) similar to that described in Low-Nam et al^29^, we estimated the dimerization kinetics between interacting receptors. In the absence of ligand, we found an off-rate (*k_off_*) for RON/RON homo-interactions of 0.18 ± 0.02 s^-1^ (average lifetime of ~5.5 sec). Together with the correlated motions analysis, these results are consistent with the idea that RON can homodimerize independent of ligand, as has been proposed by others based on the crystal structure of the RON extracellular domain^14, 29^ and the evidence for ligand-independent activation with RON overexpression or mutations in cancer^33–35^. Two-color tracking of QD655-HA-RON and QD605-EGF-EGFR returned an off-rate of 0.49 ± 0.05 s-1 for hetero-interactions. This more transient (average lifetime of ~2 sec) interaction is consistent with the ability of EGFR to phosphorylate RON without subsequent co-endocytosis.

### Maximal EGF-induced RON phosphorylation requires kinase activity of both receptors

Treatment of A431^RON^ cells with the reversible EGFR-selective kinase inhibitor, PD153035, blocks EGF-induced changes in RON mobility (Figure 3A-B). To follow up these results implicating EGFR kinase activity as the primary driver of EGFR/RON crosstalk, we treated both A431^RON^ and HEK^RON/EGFR^ cells with the irreversible pan-ErbB kinase inhibitor, afatinib. Results in Figure 5 show that afatinib treatment completely blocks EGF-dependent RON phosphorylation but does not inhibit MSP-dependent RON phosphorylation. Cells pretreated with BMS777607, a RON/Met-family kinase inhibitor, blocked MSP-dependent RON phosphorylation, but only partially blocked EGF-dependent RON phosphorylation (Figure 5B). As expected, BMS777607 did not affect EGF-dependent EGFR phosphorylation. These results indicate that both EGFR and RON kinase activity contribute to EGF-mediated RON phosphorylation.

**Figure 5.**
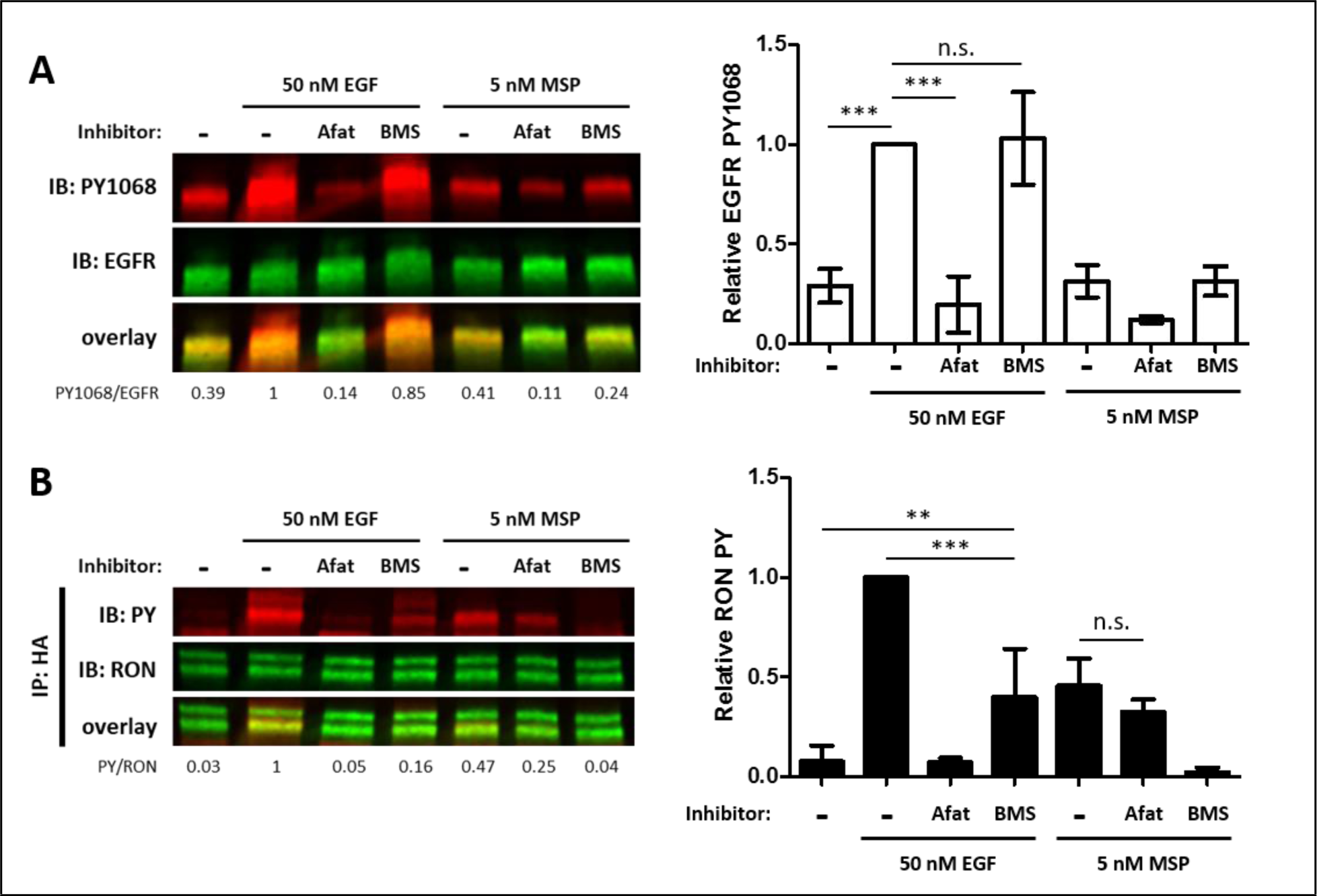
Maximal EGF-induced RON phosphrylation requires kinase activity of both receptors. (A-B) A431^RON^ cells were pre-treated with 10 μM Afatinib (Afat, pan-ErbB inhaibitor) or 1 ~M BMS777607(BMS, Met family kinase inhibitor) for 20 or 15 min, respectively. Cells were then treated ± EGF or MSP for 5 min. (A) Cell lysates were used for PY 1068 and EGFR immuniblots. (B) Lysates were immunoprecipitated (IP) with an anti-HA for PY and RON immunoblots. Bar graphs are corresponding mean ± SD from triplicate experiments. ^*^ p < 0.05; ^**^ p < 0.01; ^***^ p < 0.001.

To confirm the differential contributions of EGFR and RON kinase activity in crosstalk, we expressed the kinase dead mutant of RON (RON-K1114M) in A431 cells (**Figure S3**). EGF-driven phosphorylation of RON-K1114M was observed and afatinib treatment abrogated this phosphorylation (**Figure S3A)**. The reduction in RON phosphorylation by BMS777607, as seen in RON-WT, is not observed for RON-K1114M since this mutant lacks kinase activity. Consistent with the observed phosphorylation, A431^RON-K1114M^ cells undergo significant reduction in mobility with EGF stimulation in SPT experiments (**Figure S3B)**. These results underscore the importance of EGFR kinase activity in crosstalk and rule out Met as a possible contributor.

### EGFR/RON crosstalk does not require downstream signaling molecules

Thus far, our data indicate the critical role for EGFR kinase activity in EGF-dependent RON phosphorylation. While this could be attributed to direct phosphorylation of RON by EGFR in hetero-oligomeric complexes, an alternative mechanism could involve recruitment of EGFR-associated kinases such as the tyrosine kinase Src^36, 37^. To rule out the possibility that Src is an intermediary in propagating EGFR/RON crosstalk, A431^RON^ cells were pre-treated with the Src family kinase inhibitor dasatinib prior to stimulation with 50 nM EGF. Low doses of dasatinib (10 nM) were used to ensure Src family specificity^38^ while achieving 70% reduction in basal Src PY416 phosphorylation (**Figure S4A**). Dasatinib treatment did not inhibit EGFR kinase activity, based on detection of EGFR (PY1068), or alter RON phosphorylation (Figure 6A). Thus, EGFR/RON crosstalk does not depend on Src kinase activity.

**Figure 6.**
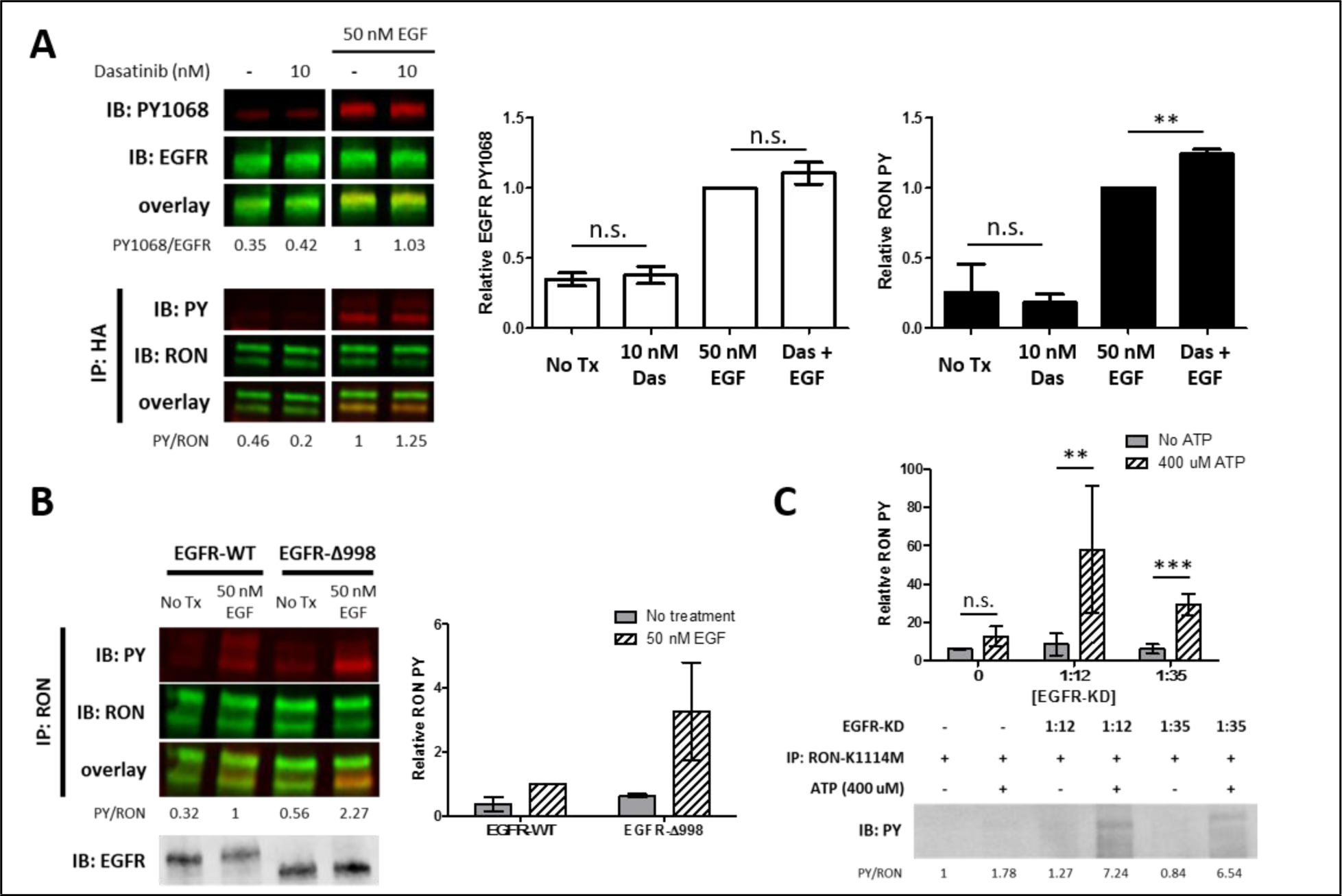
Crosstalk occurs through direct phosphorlation of RON by EGFR. (A) A431^RON^ cells were pre-treated with Dasatinib (Dnas,Src inhibitor) for 30 min prior to stimulation with EGF for 5 min at 37 °C. Representative immunoblots of cell lysates detecting PY1068 and total EGFR(top). Immunoblot detecting PY and RON after IP with anti-HA (RON). (B) HEK^RON^ cells transiently transfected with EGFR-WT or EGFR-Δ998± EGF for 5 min. Representative immunoblots detecting PY and RON after IP with anti-RON or detection of total EGFR on cell lysates (bottom inset). (C) Kinase assay using purified EGFR kinase domain (EGFR-KD) co-incubated with RON-K1114M IP samples ± ATP. Representative immunoblot detecting total phophorylation (PY) of RON. All bar graphs represent mean ± SD from triplicate experiments. ^**^ p < 0.01; ^***^ p < 0.001.

In addition to Src, EGFR also recruits a number of other cytoplasmic signaling molecules to phosphotyrosines in its C-terminal tail. We expressed in HEK^RON^ cells a version of EGFR truncated at amino acid 998 (HEK^RON/EGFR-Δ998^), which lacks most of the phosphotyrosine binding sites that recruit downstream adaptor molecules^26^. In a previous study, EGFR-Δ998 exhibited decreased phosphorylation of the remaining tyrosine residues 845, 974, and 992 compared to full length EGFR suggesting that phosphorylation at these sites might depend on downstream binding partners^26^. Unexpectedly, stimulating HEK^RON/EGFR^-Δ998 cells with EGF led to enhanced phosphorylation of RON compared to HEK ^RON/EGFR-WT^ (Figure 6B). These results confirm that recruitment of downstream signaling molecules to the C-terminal tail of EGFR is not required for EGF-driven RON phosphorylation, while raising a new question as to the mechanism of this enhanced crosstalk. We considered the possibility that truncation of the EGFR tail could prevent recruitment of EGFR-associated phosphatases that can dampen downstream signals^39, 40^. However, this possibility is unlikely, since the dephosphorylation kinetics of RON were similar in cells expressing either EGFR-WT or EGFR-Δ998 after treatment with EGF, followed by afatinib to irreversibly inhibit subsequent rounds of phosphorylation (**Figure S4B**).

### RON is a substrate for EGFR kinase activity

Having ruled out a role for downstream signaling molecules, the probable mechanism for crosstalk is that the RON C-terminal tail is a substrate for EGFR kinase activity. To test this possibility, we designed an *in vitro* kinase assay to allow for detection of EGFR phosphorylation of RON without background from other cellular components. In these experiments, we used immunoprecipitated kinase dead RON (RON-K1114M) as a substrate, removing potential contributions from RON kinase activity, and recombinant EGFR kinase domain (EGFR-KD) as the active kinase. We found that EGFR-KD directly phosphorylated RON-K1114M, in an ATP-dependent and EGFR-KD concentration-dependent manner (Figure 6C).

### RON cannot substitute as activator or receiver in EGFR dimers

Structural studies have established the critical role for the orientation of EGFR kinase domains in asymmetric dimers (activator and receiver) for EGFR kinase activity^17, 41^. We set out to determine if RON can substitute for either activator or receiver to form an active EGFR/RON heterodimer. HEK^RON^ cells were transfected with EGFR mutants that are either receiver-impaired (I682Q) or activator-impaired (V924R)^26, 42^. EGF stimulus resulted in the expected EGF-driven EGFR and RON phosphorylation patterns in HEK^RON^ cells expressing EGFR-WT (Figure 7). In contrast, neither EGFR-I682Q nor EGFR-V924R were capable of EGFR autophosphorylation or crosstalk with RON. Restoration of functional EGFR dimers by co-expression of EGFR-I682Q and EGFR-V924R rescued EGFR and RON phosphorylation. These data demonstrate that, unlike other ErbB family members that can form functional heterodimers with EGFR^43^, RON cannot serve as a substitute for the EGFR activator or receiver. Therefore, while EGFR can directly phosphorylate RON, this is not achieved through a simple heterodimerization event, but likely requires higher-order interactions (i.e. hetero-trimers or larger).

**Figure 7.**
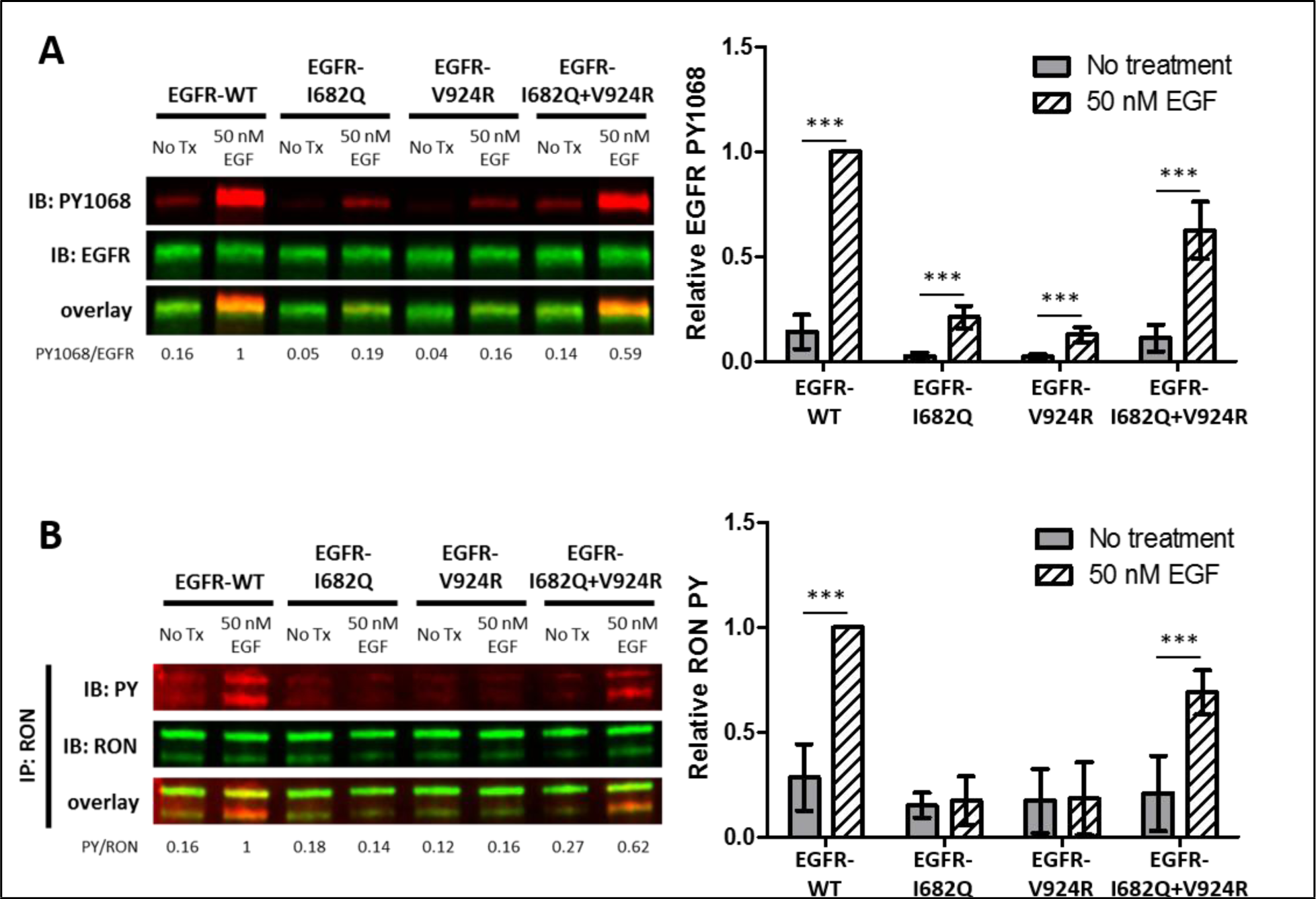
Functional EGFR dimers are necessary for EGFR/RON crosstalk. (A-B) HEK^RON^ cells transiently expressing EGFR-WT, EGFR-1682Q (receiver-impaired), EGFR-V924R (activator-impared) or both mutants (EGFR-I682Q + V924R) were treated ± EGF for 5 min at 37 ° C. (A) Representative immunoblot detecting PY 1068 and EGFR in cells lysates. (B) Representative immunoblot showing PY and RON after IP with anti-RON. Triplicate experiments from A and B are quantified and graphed as mean ± SD. ^***^ p < 0.001.

## DISCUSSION

A key step in RTK signaling is the ligand-driven formation or rearrangement of homo- and hetero-oligomers that enable auto- and trans-activation of the cytoplasmic kinase domains^29, 44–47^. While previous studies have demonstrated crosstalk between EGFR and RON, the molecular mechanisms that facilitate this signaling outcome are not known. Our studies revealed that crosstalk occurs through direct receptor interaction and that RON is a substrate for the EGFR kinase domain. We also provide new insight into the structural requirements for productive RON and EGFR interaction. We show that RON cannot substitute as an activator or receiver kinase domain with EGFR. Therefore, transactivation of RON by EGFR first requires formation of a signaling competent EGFR dimer.

This work shows definitively that crosstalk is EGF-driven and propagates in a unidirectional manner from EGFR to RON. Others have suggested that EGFR and RON can transactivate each other^6, 12^. One explanation for the previous findings could be cross-reactivity of anti-phosphotyrosine antibodies used, since we found some “receptor-specific” commercial antibodies to be cross-reactive with phospho-RON and phospho-EGFR. To address this potential problem, we carefully validated the antibodies used in our experiments (see **Figure S1A**) and confirmed that our SDS-PAGE methods effectively resolved the RON pro-form from mature RON for western blotting protocols. We also considered the possibility that crosstalk could be dependent on the ratio of EGFR/RON levels, developing cell lines where EGFR is highly overexpressed compared to RON (~24:1) or where the expression is similar (~2:1). In both cases, crosstalk was found to be unidirectional and EGF-dependent. Finally, to focus on interactions between wild type EGFR and wild type RON, we selected model cell lines lacking the endogenous expression of other RON splice variants. Future studies are needed to define the role of crosstalk in situations where RON is more abundant than EGFR or different isoforms of RON are present.

Unidirectional crosstalk has been reported for other RTK combinations involving EGFR. For example, EGFR was found to transactivate Met, but not vice versa, in human hepatoma and human epidermal carcinoma (A431) cell lines^48^. van Lengerich et al. described unidirectional receptor phosphorylation between EGFR and ErbB3, where the addition of EGF leads to ErbB3 phosphorylation, but there is no change in EGFR phosphorylation in response to stimulation with ErbB3’s ligand, neuregulin49. They suggested a mechanistic model whereby activated EGFR homodimers engage in higher order oligomers with ErbB3 precluding the need for ErbB3 to function as an allosteric activator of EGFR.

We explored these same concepts with respect to EGFR/RON crosstalk. Our co-precipitation studies support a model of direct interaction, based upon both western blotting and mass spectrometry read-outs. Consistent with other reports^6, 12^, the ability to co-IP EGFR with RON was independent of ligand addition. Ligand-independent interactions have similarly been reported for RON with Met5 and Met with EGFR^48^. Based upon results from immunoelectron microscopy, resting EGFR and RON are located in close proximity on the cell surface. Direct interaction between EGF-bound EGFR and RON were observed in our two-color SPT experiments. It is noteworthy that, although complexes containing EGF-EGFR and RON are stable enough to promote RON phosphorylation, they disengage prior to EGFR endocytosis. This is consistent with previous studies where EGFR and ErbB3 did not co-endocytose after EGF stimulation^32^.

Others have found that EGFR/Met crosstalk in the presence of the EGFR inhibitor gefitinib can be mediated by Met phosphorylation of the cytoplasmic tyrosine kinase c-Src^50^. The cytoplasmic tail of EGFR contains multiple tyrosines that are known to recruit adaptor proteins when phosphorylated, including Grb2, Shc and c-Src^51, 52^. Our use of adaptor protein-specific inhibitors and structural mutants ruled out this mechanism for EGFR/RON crosstalk. We found that neither inhibition of c-Src activity nor removal of the EGFR cytoplasmic tail (EGFR-Δ998) prevented crosstalk to RON. In fact, expression of EGFR-Δ998, which lacks the majority of EGFR phosphorylation sites, but retains an autoinhibitory domain^26^, resulted in enhanced RON phosphorylation upon EGF treatment. This increased activation was not due to the lack of phosphatase recruitment, as dephosphorylation rates of RON were the same in the presence of either wildtype or truncated EGFR. It is possible that other signaling molecules play a negative regulatory role. Grb2 has been reported to inhibit RON autophosphorylations^53^, raising the possibility that the loss of Grb2 recruitment by EGFR-Δ998 would reduce local Grb2 concentration and increase RON phosphorylation. A more probable interpretation is that removal of the C-terminal tail eliminates transphosphorylation of the EGFR tail itself, and diminishes the recruitment of downstream EGFR substrates, so that the RON C-terminal tail becomes the preferred substrate in the hetero-oligomeric complexes. Thus, while a third-party signaling molecule is not required to mediate crosstalk in our model systems, the unstructured EGFR tail or its binding partners appear to have a role in limiting EGFR-mediated phosphorylation of RON.

The structural requirements for direct interactions between EGFR and RON are yet unresolved. Considering the simplest model, we tested whether RON can act as an activator or receiver kinase in an EGFR/RON dimer. *In vitro* kinase assays demonstrated that RON is indeed a substrate for the EGFR kinase. However, expression of either receiver-impaired (I682Q) or activator-impaired (V924R) EGFR mutants could not support crosstalk with RON. Co-expression of EGFR-I682Q and EGFR-V924R that restored EGFR autophosphorylation also restored crosstalk. This indicates that RON and EGFR do not interact to form simple heterodimers. Instead, an signaling-competent EGFR homodimer appears to be required for EGF-driven RON phosphorylation. Considering that RON homo-interactions were observed by two-color SPT, in both resting and liganded states, we postulate that the complex is minimally a dimer of dimers (tetramer), but further study is needed to determine this exact stoichiometry.

Our results highlight a potential therapeutic escape mechanism for RON-driven oncogenesis, in that RON signaling outcomes can be stimulated by EGFR in the absence of MSP and even if RON kinase activity is inhibited. This may be one explanation for why a phase I clinical trial for the anti-RON neutralizing antibody, narnatumab, did not show significant antitumor activity as a monotherapy^54^. In support of dual therapy, studies in mice showed that neutralizing antibodies against RON exhibit 50% decreased pancreatic xenograft tumor growth that is enhanced in combination with an EGFR inhibitor^55^. The novel molecular mechanisms governing EGFR/RON crosstalk described here have provided new insights into therapeutic resistance and suggest that disruption of interactions between RON and EGFR could provide a therapeutic advantage.

## ACKNOWLEDGEMENTS

This work was supported by the National Institutes of Health (NIH) R35GM126934 (DSL), NIH R21 GM132716 (KAL) and the New Mexico Spatiotemporal Modeling Center (NIH P50GM085273). Additional support was from the UNM MARC Program (NIH 2T34GM008751) to JMK, the ASERT-IRACDA Program (NIH K12GM088021) to EDJ, and the UNM Undergraduate Pipeline Network to JMK and ACG. We thank Peter Cooke of the Electron Microscopy Laboratory at New Mexico State University for access to a Hitachi H-7650 TEM. Mass spectrometry data was generated at the UT Southwestern Proteomics Core. We thank Shayna Lucero for assistance with cell culture and flow cytometry, Dr. Michael Wester for assistance with EM analysis, Mohamad Fazel for assistance with single particle tracking analysis, Dr. Chris Valley and Russell Hunter with plasmid preparations and Dr. Mark Lemmon for helpful discussions. We gratefully acknowledge use of the University of New Mexico Comprehensive Cancer Center fluorescence microscopy and flow cytometry facilities, as well as the NIH P30CA118100 support for these cores.

## Author Contributions

Plasmids were generated by EWG, CFN, and MPS. EWG generated the A431^RON^ cell line and performed initial biochemical and SPT experiments. RMG prepared, acquired, and analyzed the Electron Microscopy images. CFN performed all biochemical assays with support from AG. CFN performed the flow cytometry cell receptor characterization and generated the HEK^RON^ and HEK^RON/EGFR^ cell lines. CFN and JMK performed the immunofluorescence experiments. EDJ and IO-C performed SPT experiments and diffusion analysis. KAL and DJS performed two-state HMM analysis. DSL directed the project. CFN, EWG, AR, MPS, BSW, and DSL participated in the conceptualization and design of the work. CFN, MPS, BSW, and DSL designed and interpreted experiments, and wrote the manuscript with input from all authors.

## METHODS

### Cell lines and Reagents

Cell culture medium was from Thermo Fisher Scientific and Poly-L-lysine (PLL) from Sigma (cat # P4707). Afatinib and BMS777607 were from Selleck Chemicals (cat # S1011 and S1561, respectively), Dasatinib from Santa Cruz Biotechnology (cat # sc-358114), and PD153035 from EMD Millipore (cat # 234491). Human recombinant EGF was from Invitrogen (cat # PHG0311) or PeproTech (cat # AF-100-15), biotin-conjugated and AF647-conjugated EGF from Thermo Fisher Scientific (cat # E3477 and E35351), and MSP from R&D Systems (cat # 4306-MS-010). Halt protease and phosphatase inhibitor (PPI) cocktail was from Pierce (cat # 78446) and the protease inhibitor cocktail set V, EDTA-free was from Calbiochem (cat # 539137). Quantum dot (QD) 605 and QD 655 streptavidin conjugates were from Thermo Fisher Scientific (cat # Q10101MP and Q10121MP, respectively). For western blotting, BCA protein assay kit (cat # 23225) and ECL blotting substrate (cat # 32106) were from Pierce. Immunoprecipitation was based on use of protein A/G magnetic beads from Pierce (cat # 88802). See **Table S2** for a list of primary and secondary antibodies used in these studies.

Human epidermoid carcinoma A431 cells (ATCC, CRL-1555) were cultured in Dulbecco’s Modified Eagle Medium (DMEM) supplemented with 10% HyClone cosmic calf serum (CCS; GE Healthcare Life Sciences), 2 mM L-glutamine (Life Technologies), and penicillin/streptomycin (Life Technologies). Human embryonic kidney HEK-293 cells were cultured in Minimum Essential Medium (MEM) with 10% fetal bovine serum (FBS; Atlanta Biologicals), 2 mM L-glutamine, and penicillin/streptomycin.

### Plasmid cloning, site directed mutagenesis and cell transfections

The vector containing RON (MST1R) pDONR223-MST1R was a gift from William Hahn and David Root (Addgene plasmid # 23942; http://n2t.net/addgene:23942;RRID:Addgene_23942)^56^. HA-tagged RON was cloned into the expression vector pcDNA3.1/V5-His-TOPO (Invitrogen) by fusion PCR. An ultramer containing the CACC ligation sequence, start codon, RON signal peptide, HA-tag, and alanine linker 5’ of the mature RON coding region and a reverse primer were used to synthesize HA-RON. DNA oligoes were from Integrated DNA Technologies. Ultramer sequencing and mutagenesis primers are listed in **Table S3**. The kinase dead RON variant (HA-RON-K1114M) was generated by site-directed mutagenesis36 (**Table S3**). To establish cell lines stably expressing HA-RON (HEK^RON^ and A431^RON^), cells were transfected with the pcDNA3.1 HA-RON plasmid by electroporation using the AMAXA Nucleofector System (Lonza). Briefly, 5 × 10^6^ HEK-293 cells were transfected with 8 µg of plasmid DNA using Nucleofection Solution V and program Q-001. A431 cells were transfected with HA-RON or HA-RON-K1114M using solution T and program X-001. Transfected cells were selected for stable integration by growth in 1 mg/ml G418 (Caisson Labs) for 7 days, then sorted for RON expression with a fluorescently-conjugated anti-HA antibody using an iCyt SY3200 cell sorter (Sony Biotechnology).

For co-expression of RON and EGFR, HEK^RON^ cells were transfected with an ACP-tagged EGFR plasmid^28^ by electroporation using the same conditions as above. Transfected cells were selected with zeocin (300 ug/ml; Gibco/Life Technologies) and sorted for double positive cells (anti-HA-AF488 and anti-EGFR-AF647) on the iCyt SY3200.

For kinase assays, a C-terminal SBP tagged construct of EGFR encoding the transmembrane and kinase domain (EGFR-KD) was amplified from full length EGFR by PCR (**Table S3**) and cloned into the pCTAP backbone via Gibson assembly. EGFR-KD and RON-K1114M proteins were produced using the Expi293 cell Expression System (Thermo Fisher Scientific) according to the manufacturer’s recommendations.

Receiver-only and activator-only EGFR mutants, EGFR-I682Q and EGFR-V924R, were engineered from the pcDNA3.1 HA-EGFR WT plasmid using site-directed mutagenesis^28^ (**Table S3**). The truncated EGFR-Δ998 plasmid, which lacks the C-terminal phosphorylation sites, was generated by amplifying the truncated EGFR from pcDNA3.1-EGFR WT plasmid using standard PCR and cloning techniques (**Table S3**). HEK^RON^ cells were transiently transfected with the EGFR mutants and experiments performed at ^18–24^ hr post-transfection.

### Flow cytometry – receptor quantification

Cell surface expression quantification of EGFR and RON was analyzed by flow cytometry using Quantum MESF kits. Briefly, cells were incubated with a range of concentrations (0-40 ug/ml) of anti-EGFR-AF647 (dye/protein ratio of 2.74 or 3.84) or anti-HA-AF488 (dye/protein ratio of 3.34) for 1 hr on ice. Cells were rinsed with PBS, fixed in 4% PFA (paraformaldehyde) for 10 min on ice, washed with 10 mM Tris-PBS and resuspended in PBS. Fluorescent calibrator beads, Quantum AlexaFluor 647 or 488 MESF (Bangs Laboratories, cat # 647A and 488A, respectively) were used to generate a standard curve of fluorescence intensity. Samples and beads were run on the Accuri C6 Plus cytometer (BD Biosciences), and receptor levels calculated based on the dye:protein ratio of the individual antibodies and values determined using the QuickCal spreadsheet (Bangs Laboratories).

### Immunofluorescence Staining

HEK^RON/EGFR^ cells were plated onto glow-discharged (EMS 150 T ES, Quorum Technologies), PLL coated glass coverslips overnight. RON labelling was done in live cells with an anti-HA-FITC fragment antibody (Fab) for 30 min in Tyrodes buffer (135 mM NaCl, 10 mM KCl, 0.4 mM MgCl^2, 1^ mM CaCl2, 10 mM HEPES, 20 mM glucose, 0.1% BSA, pH 7.2) on ice. Cells were treated with 10 nM EGF-AF647 on ice for 5 min, fixed in 4% PFA for 15 min at RT, and washed with 10 mM Tris / PBS buffer. Samples were rinsed, incubated with DAPI, and mounted with Prolong Gold or Diamond (Thermo Fisher Scientific). Confocal images were acquired using a 63x/1.40 oil objective on a Zeiss LSM800 microscope in channel mode and appropriate diode lasers were used for excitation of the fluorophores.

For endocytosis experiments, RON was pre-labeled with anti-HA-FITC Fab and cells were stimulated with EGF-AF647 for 10 min at 37°C prior to fixation. Samples were simultaneously blocked and permeabilized with 0.1% Triton X-100 / 3% BSA / PBS for 20 min and stained with anti-EEA1 antibody in 0.1% Triton X-100 / 0.1% BSA / PBS solution for 30 min at 37°C followed by anti-Rabbit-AF555 secondary for 30 min at 37°C before DAPI staining and mounting.

### Transmission Electron Microscopy of Native Membrane Sheets

Standard “rip-flip” membrane sheets were prepared as previously described^57^. In brief, A431^RON^ cells were treated or not with 50 nM EGF for 2 or 5 min and fixed in 0.5% PFA. Coverslips were flipped, cells down, onto PLL coated formvar and carbon-coated nickel finder grids and pressure was applied to adhere apical cell membranes before removing the coverslip. Grids with membrane sheets were fixed with 2% PFA in HEPES buffer (25 mM HEPES, 25 mM KCl, and 2.5 mM Mg Acetate) for 20 min and sequentially labeled with anti-RON or anti-EGFR antibodies in 0.1% BSA / PBS for 1 hr at RT. Secondary antibodies conjugated to colloidal gold were added for 30 min at RT. Samples were post-fixed with 2% glutaraldehyde for 20 min and negatively stained with 0.3% tannic acid for 1 min and 2% uranyl acetate for 9 min. Digital images were acquired on a Hitachi H-7650 Transmission Electron Microscope equipped with a mid-mount digital imaging system (Advanced Microscopy Techniques, Corp) and Image J (NIH) was used to crop images. Ripley's bivariate K test was used to determine if co-clustering of species is significant^23, 24^. Data within the confidence window is not co-clustered, while data plotted above or below the line is found to be significant, either hetero-or homo-clustered, respectively.

### Cell activation and lysis

Transiently transfected or stable cell lines were seeded into 100 mm dishes and allowed to adhere overnight. For inhibition studies, cells were pretreated with 10 μM Afatinib for 20 min, 1 μM BMS777607 for 15 min, or ^1–10^ nM Dasatinib for 30 min, where indicated. They were subsequently treated with different doses of EGF, MSP, or both, for varying times (0 to 5 min). Cells were rinsed in cold PBS and lysed on ice for 20 min with NP-40 lysis buffer (150 nM NaCl, 50 mM Tris, 1% NP-40) containing PPI. Lysates were cleared and protein concentrations in the supernatant was determined by BCA protein assay.

### Immunoprecipitation

Cell lysates (1 mg total protein) were immunoprecipitated (IP) overnight using an anti-HA antibody coupled to magnetic or sepharose beads or anti-RON antibody overnight at 4°C, rotating. For samples incubated with the anti-RON antibody, protein A/G magnetic beads were added the next day and incubated for 1h, rotating at 4°C. Beads were washed with 0.05% Tween-20 / PBS containing PPI.

### Multiplex Immunoblotting

Whole lysates (20 ug) or IP samples were boiled with reducing sample buffer, subject to SDS-PAGE, and transferred to nitrocellulose membranes using the iBlot2 system (Life Technologies). Membranes were blocked for 30 min in 3% BSA / 0.1% Tween-20 / TBS, and probed overnight with primary antibodies at 4°C (**Table S2**). Membranes were incubated with IRDye fluorescent secondary antibodies for one hr at RT (**Table S2**), washed, and dual color detection was performed using the Odyssey Fc Imaging System (Li-Cor). Band intensities were analyzed with Image Studio (Li-Cor, version 5.2) and normalized PY to total protein (PY1068/EGFR or PY/RON).

### Single Particle Tracking (SPT)

Single- and dual-color SPT and analysis was conducted as previously described^28–30^. Briefly, A431^RON^ (Figure 3A-B and 4) or A431^RON-K1114M^ (**Figure S3B**) cells were seeded in 8-well chamber slides (Nunc Lab-Tek) at a density of 30,000/well and allowed to adhere overnight. Where indicated, EGFR kinase activity was inhibited by pretreating with 1 μM PD153035 for 2 hr and maintained throughout the experiment. RON was tracked via QD-conjugated to biotinylated anti-HA Fab fragments that bind to the N-terminal HA-tag on HA-RON (as indicated). Cells were incubated with 200 pM anti-HA-QDs (605 or 655) for 15 min at 37°C to obtain single-molecule density on the apical surface. After washing with Tyrodes buffer cells were treated with 5 nM MSP for 5 min or 50 nM EGF for 30 sec and imaged. For dual EGFR and RON tracking, cells were incubated with 200 pM anti-HA-QD655 for 15 min at 37°C, washed, and then stimulated with 50 pM QD605-conjugated EGF-biotin. Particle tracking was done for up to 15 min. Imaging was performed on an Olympus IX71 inverted wide field microscope with a 60x 1.2 numerical aperture water objective as in Valley *et al*, 2015^28^. QD emissions were collected using a 600 nm dichroic (Chroma) and the appropriate bandpass filters, 600/52 nm and 676/37 (Semrock). Physiological temperature (34 - 36°C) was maintained using an objective heater (Bioptechs). Images were acquired at a rate of 20 frames per sec for a total movie length of 1,000 frames.

MATLAB (MathWorks) was used for image processing and analysis in conjunction with DIPImage (Delft University of Technology). Diffusion was computed using mean square displacement (MSD)^28, 29, 58^. Dimer off-rates and events were identified using a two-state HMM, similar to previous work^29, 30^. For more details see Supplemental Methods.

### Protein purification and Kinase assay

EGFR-KD from Expi293 cell lysates was bound to streptavidin resin and eluted in biotin buffer according to manufacturer’s recommendations (InterPlay Mammalian TAP System; Agilent Technologies). Typical protein yield was between 100-400 ng/µl. RON-K1114M was immunoprecipitated from Expi293 cell lysates (2 mg total protein) with sepharose anti-HA beads. Immunoprecipitated RON-K1114M was resuspended in kinase assay buffer (200 mM HEPES, pH 7.4; 300 mM MgCl_2_; 20 mM MnCl_2_; 0.5% Triton X-100; 1.5% Brij 35; 10% glycerol; 1X Protease inhibitor cocktail Set V; and 2 mM activated Na_3_VO_4_) in the presence or absence of purified EGFR-KD (1:12 or 1:35 dilution). Samples were incubated with 400 µM ATP (or no ATP, as a control; Cell Signaling Technology, cat # 9804) and held at 30°C for 30 min, shaking. Reactions were terminated by addition of ice cold buffer, RON-K1114M bound to beads was recovered by centrifugation at 2500 x g for 2 min at 4°C and washed 3× with 0.05% Tween-20 / PBS containing PPI. Samples were boiled with reducing sample buffer, subject to SDS-PAGE, and western blotting with HRP-conjugated anti-PY20 and anti-PY99 antibodies.

### Statistical Analysis

All values from quantitative western blot experiments are plotted as mean ± SD. For quantitative experiments, statistical analysis was performed using GraphPad Prism (Prism 4, GraphPad) with a two-way analysis of variance (ANOVA) from three separate replicate experiments. For immunoblot analysis, phosphorylated protein levels were normalized to total protein levels (RON or EGFR) detected in the same samples. For phosphorylation time course experiments, the maximum stimulation level was set at one for triplicate experiments. Differences among means were tested using the Bonferroni multiple comparison test. Values of p < 0.05 were considered significant. Errors in values of diffusion coefficients are reported as 95% confidence intervals from fitting a Brownian diffusion model (linear) to the first 5 points of the MSD.

## VIDEO LEGENDS

**Video 1.** Two RON receptors are engaged in an interaction from the start of the video, which lasts for ~ 5 sec before the receptors dissociate. This movie accompanies the interaction shown in Figure 4B. QD605 is displayed in green and QD655 in magenta. A colored tail for each QD shows a track of the previous 10 localizations. Playback speed is 20 frames/sec. Images have been brightness and contrasts enhanced for visualization.

**Video 2.** Two RON receptors undergoing repeated interactions that each last ^1–2^ sec. This movie accompanies the interaction shown in **Figure S2A**. Color scheme, comet tail, and payback speed are as previously mentioned for Video 1.

**Video 3.** A long-lived RON/RON interaction. This movie accompanies the interaction shown in **Figure S2B**. Color scheme, comet tail, and payback speed are as previously mentioned for Video 1

**Video 4.** A short-lived interaction between QD605-EGF-EGFR (green) and QD655-RON (magenta). This movie accompanies the interaction shown in Figure 4E. A colored tail for each QD shows a track of the previous 10 localizations. Playback speed is 20 frames/sec. Images have been brightness and contrasts enhanced for visualization.

**Video 5.** A complex of EGF-EGFR and RON is seen to break apart after a ~ 3.5 sec dimer event. This movie accompanies the interaction shown in **Figure S2C**. Color scheme, comet tail, and payback speed are as previously mentioned for Video 4.

**Video 6.** A long-lived interaction between EGF-EGFR and RON. This movie accompanies the interaction shown in **Figure S2D**. Color scheme, comet tail, and payback speed are as previously mentioned for Video 4.

