## Supplemental Figures, Tables, and Methods for "EGFR transactivates RON to drive oncogenic crosstalk"

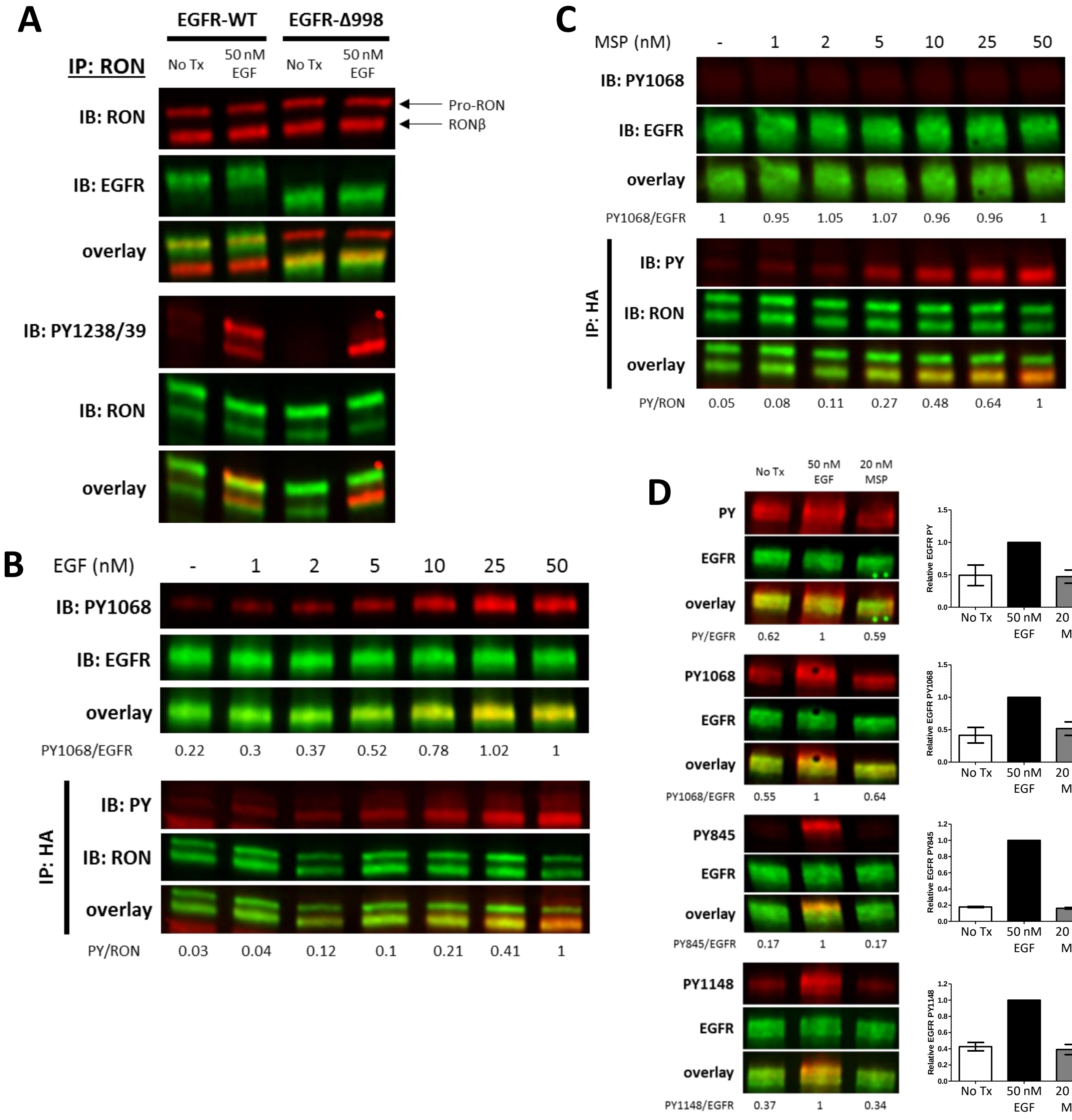


**Supplementary Figure 1. Antibody selectivity and characterization of cell models.**
(A) HEK$^{RON}$ cells transiently transfected with EGFR-WT or EGFR-Δ998 ± EGF treatment for 5 min. Cell lysates were immunoprecipitated (IP) with anti-RON and probed for EGFR, RON, or RON PY1238/39. Arrows highlight the location of the 180 kDa unprocessed RON (pro-RON), and the 145 kDa RONβ band (mature RON). Truncated EGFR-Δ998 appears as a lower molecular weight band in the anti-EGFR immunoblot, as expected. However EGFR-WT co-migrates with unprocessed RON (180 kDa). Therefore, all experiments here quantify the bottom RON band (RONβ). Additionally, the top band in the PY1238/39 immunoblot for EGF-treated EGFR-WT cells is absent in the EGFR-Δ998 samples, confirming this antibody cross-reacts with phosphorylated EGFR. Experiments to quantify RON phosphorylation were therefore first IP for HA (RON), and then probed with PY antibodies.
(B and C), A431$^{RON}$ cells were treated with increasing levels of EGF (B) or MSP (C) for 5 min. Representative immunoblots demonstrating detection of PY1068 and EGFR on cell lysates or detection of PY and RON on samples IP with anti-HA (RON).
(D) HEK$^{RON/EGFR}$ cells were treated ± EGF or MSP for 5 min. Representative immunoblots demonstrating detection of total PY (PY20/PY99 cocktail), PY1068, PY845, PY1148, and EGFR on cell lysates.

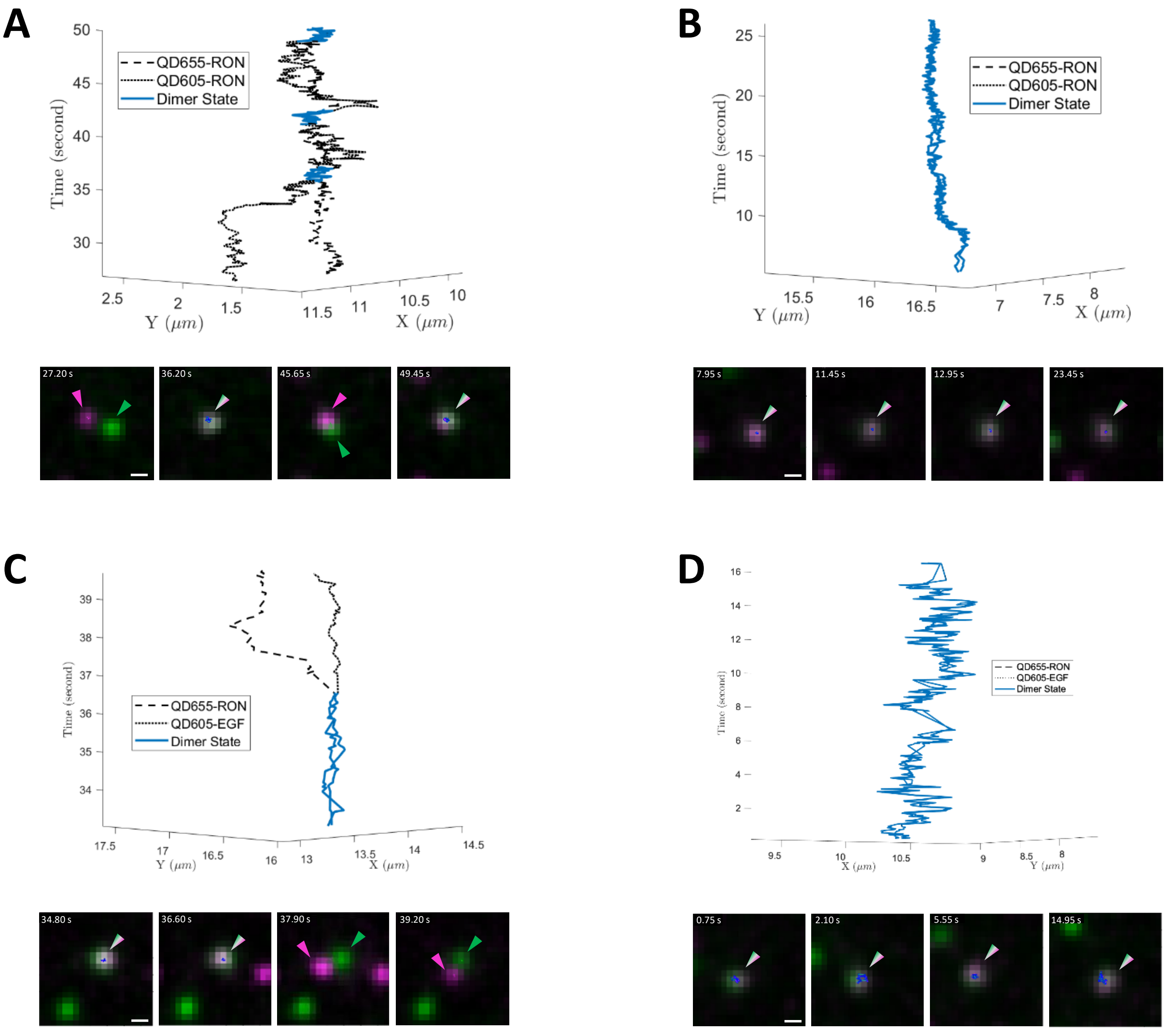


**Supplementary Figure 2. Additional examples of RON/RON and EGFR/RON interactions by two-color QD tracking.**

(A) Example 3D trajectory (top) of two QD-RON receptors that engaged in repeated transient interactions (blue segments).

(B) Example 3D trajectory (top) of a long-lived (~22 s) interaction between two QD-RON receptors. Still images from each time series are shown below the 3D trajectory. Scale bar, 500 nm.

(C) Example 3D trajectory (top) of EGF-bound EGFR and RON receptors that are initially found in a dimer complex (blue) that then dissociates at 36.5 s.

(D) Example 3D trajectory (top) of a long-lived (~16 s) interaction between two QD-RON receptors. Still images from each time series are shown below the 3D trajectory. Scale bar, 500 nm.

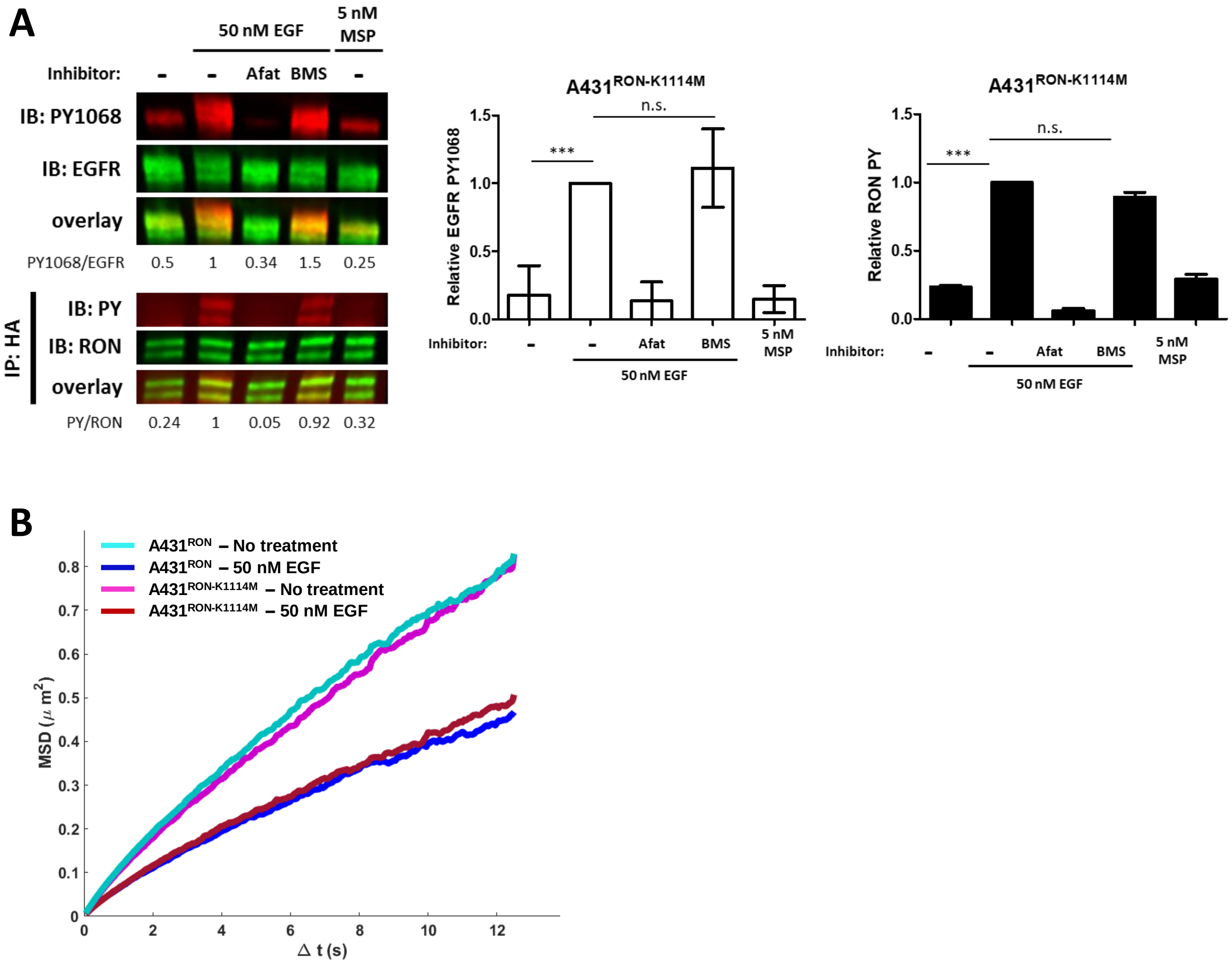


**Supplementary Figure 3. Kinase dead RON confirms EGF-induced RON phosphorylation requires kinase activity of both receptors.**

(A) A431$^{RON-K1114M}$ (RON kinase dead) cells were pre-treated, where indicated, with Afatinib (Afat; pan-ErbB inhibitor) or BMS777607 (BMS; Met family kinase inhibitor) for 20 or 15 min, respectively. Cells were subsequently treated with EGF or MSP for 5 min and lysed for immunoblot analysis. For RON and phospho-RON immunoblotting, lysates were IP with an anti-HA antibody (RON) and whole cell lysates used for EGFR and EGFR-PY1068, and their respective overlays. Triplicate experiments for EGFR (white bars) and RON (black bars) phosphorylation are plotted as mean ± SD. *** $p < 0.001$.

(B) A431$^{RON}$ or A431$^{RON-K1114M}$ cells were treated ± EGF for 5 min before imaging. RON diffusion was tracked using QD605-HA-RON and measured over time, displayed as an ensemble mean squared displacement (MSD) plot. A reduction in slope of the MSD indicates a reduces mobility.

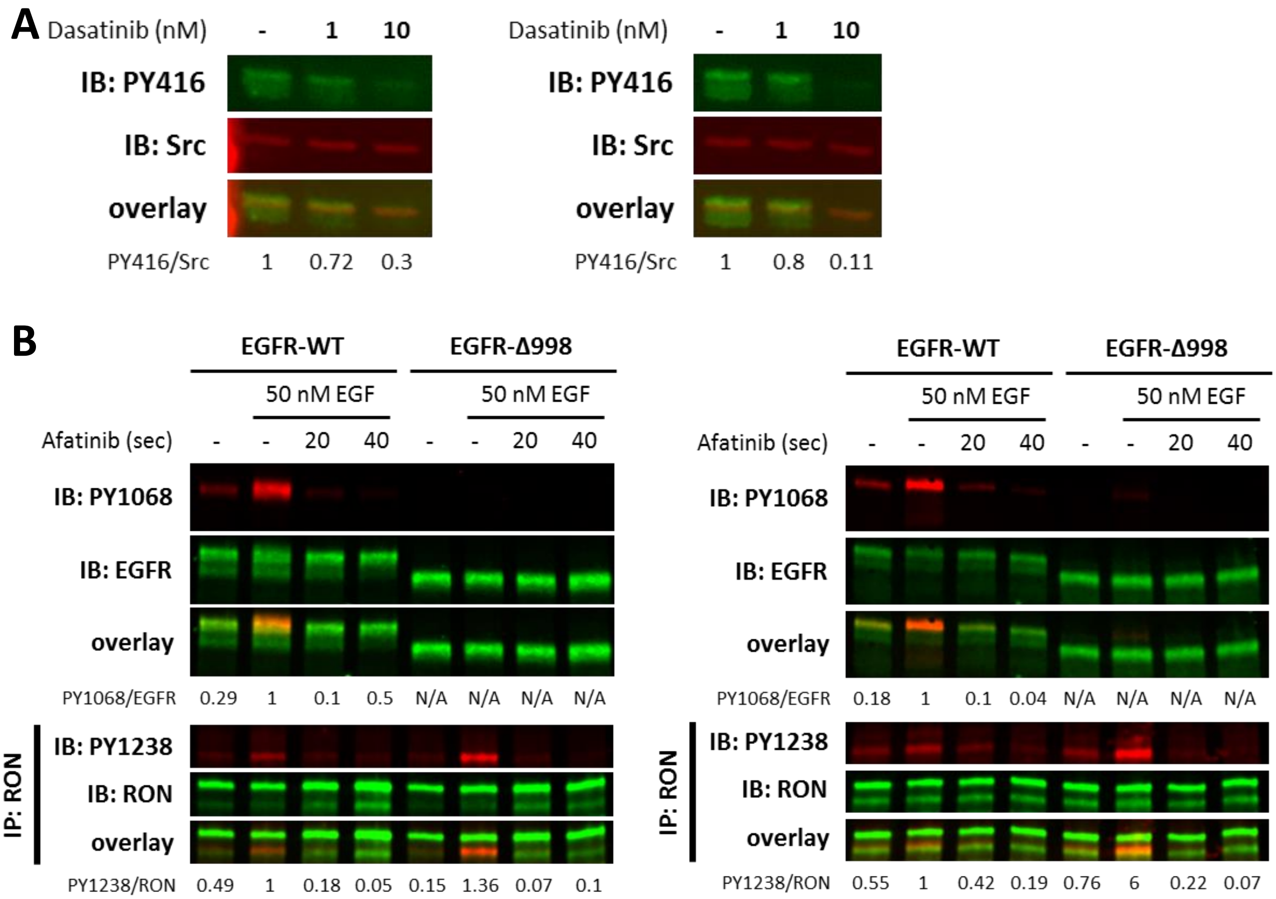


**Supplementary Figure 4. EGFR/RON crosstalk is independent of downstream signaling molecules.**
(A) A431$^{RON}$ cells were pre-treated with Dasatinib (Src inhibitor) at different concentrations for 30 min prior to cell lysis. Cell lysates were probed by immunoblot for Src-PY416 and total Src, with corresponding overlays. Immunoblots show repeat experiments.
(B) Dephosphorylation assay was conducted using HEK$^{RON}$ cells transiently transfected with EGFR-WT or EGFR-Δ998. Cells were activated ± EGF for 2 min, and immediately treated with 10 μM Afatinib (pan-ErbB inhibitor) for 20 or 40 sec. Two replicate experiments show detection of EGFR-PY1068 and EGFR on cell lysates or PY1238/39 and RON on samples IP with anti-RON, and their respective overlays.

#### SUPPLEMENTARY TABLES

**Supplementary Table S1.** List of top proteins co-IP in A431$^{RON}$ cells with anti-HA antibody for RON pulldown, as analyzed by Mass Spectrometry.

| Gene | Description (UniProt Accession) | Molecular Weight **(KDa)** | **Peptide Spectrum Matches** | **Sequence Coverage (%)** | **Spectral counts: No Tx** | **Spectral counts: 50 nM EGF** | **Spectral counts: 5 nM MSP** |
|---|---|---|---|---|---|---|---|
| **RON** | Macrophage-stimulating protein receptor (**Q04912**) | 152.6 | 1474 | 83.40 | 458.85 | 433.40 | 404.15 |
| **RPB2** | DNA-directed RNA polymerase II subunit RPB2 (**P30876**) | 134.2 | 604 | 67.20 | 227.00 | 137.00 | 188.00 |
| **RRP12** | RRP12-like protein (**Q5JTH9**) | 144.0 | 538 | 78.70 | 176.00 | 131.00 | 184.00 |
| **MTCL1** | Microtubule cross-linking factor 1 (**Q9Y4B5**) | 210.0 | 444 | 74.60 | 153.00 | 101.00 | 153.00 |
| **SOGA1** | Protein SOGA1 (**O94964**) | 160.1 | 429 | 77.90 | 136.00 | 97.00 | 152.00 |
| **MA7D1** | MAP7 domain-containing protein 1 (**Q3KQU3**) | 93.0 | 390 | 78.00 | 162.00 | 59.00 | 125.00 |
| **DLG5** | Disks large homolog 5 (**Q8TDM6**) | 214.3 | 358 | 67.70 | 127.00 | 75.00 | 138.00 |
| **BCAR1** | Breast cancer anti-estrogen resistance protein 1 (**P56945**) | 93.6 | 328 | 88.00 | 115.50 | 63.50 | 118.50 |
| **EGFR** | Epidermal growth factor receptor (**P00533**) | 134.6 | 302 | 69.30 | 98.53 | 74.04 | 86.04 |
| **RECQ4** | ATP-dependent DNA helicase Q4 (**O94761**) | 133.4 | 296 | 70.40 | 95.00 | 81.00 | 99.00 |
| **PHLB2** | Pleckstrin homology-like domain family B member 2 (**Q86SQ0**) | 142.4 | 283 | 74.10 | 108.50 | 51.50 | 96.50 |
| **CE170** | Centrosomal protein of 170 kDa (**Q5SW79**) | 175.7 | 279 | 70.90 | 106.00 | 37.50 | 97.50 |
| **K2C1** | Keratin, type II cytoskeletal 1 (**P04264**) | 66.2 | 264 | 76.70 | 52.50 | 115.50 | 66.50 |
| **RHG23** | Rho GTPase-activating protein 23 (**Q9P227**) | 162.5 | 253 | 69.40 | 90.50 | 58.50 | 87.00 |
| **UACA** | Uveal autoantigen with coiled-coil domains and ankyrin repeats (**Q9BZF9**) | 162.8 | 253 | 68.10 | 95.58 | 25.58 | 103.58 |
| **KIF14** | Kinesin-like protein KIF14 (**Q15058**) | 186.9 | 253 | 62.40 | 91.00 | 57.00 | 95.00 |
| **PLEC** | Plectin (**Q15149**) | 532.9 | 251 | 32.90 | 101.00 | 24.50 | 106.50 |
| **PKHA7** | Pleckstrin homology domain-containing family A member 7 (**Q6IQ23**) | 127.4 | 245 | 73.40 | 94.00 | 51.00 | 81.00 |
| **LAMC2** | Laminin subunit gamma-2 (**Q13753**) | 131.2 | 242 | 65.80 | 81.00 | 71.00 | 79.00 |
| **PKHA5** | Pleckstrin homology domain-containing family A member 5 (**Q9HAU0**) | 127.7 | 241 | 79.40 | 95.00 | 52.00 | 81.00 |
| **RHG32** | Rho GTPase-activating protein 32 (**A7KAX9**) | 231.0 | 241 | 61.10 | 96.99 | 39.99 | 87.00 |
| **A0A0U1RQF3** | Uncharacterized protein (**A0A0U1RQF3**) | 118.9 | 233 | 77.00 | 82.00 | 55.00 | 80.00 |

| | | | | | | | |
|---|---|---|---|---|---|---|---|
| **CLAP2** | CLIP-associating protein 2 (**O75122**) | 141.4 | 226 | 67.20 | 77.50 | 52.50 | 72.50 |
| **PTN14** | Tyrosine-protein phosphatase non-receptor type 14 (**Q15678**) | 135.5 | 221 | 59.10 | 79.50 | 46.50 | 71.50 |
| **K1C9** | Keratin, type I cytoskeletal 9 (**P35527**) | 62.2 | 221 | 96.60 | 48.00 | 95.00 | 56.00 |
| **PKCB1** | Protein kinase C-binding protein 1 (**Q9ULU4**) | 132.0 | 218 | 60.20 | 81.00 | 42.00 | 80.00 |
| **KIF7** | Kinesin-like protein KIF7 (**Q2M1P5**) | 150.9 | 218 | 65.30 | 84.99 | 48.99 | 77.99 |
| **CLH1** | Clathrin heavy chain 1 (**Q00610**) | 192.0 | 212 | 64.50 | 93.00 | 29.00 | 75.00 |
| **SHRM3** | Protein Shroom3 (**Q8TF72**) | 217.3 | 210 | 54.50 | 82.00 | 39.00 | 69.00 |
| **CAR10** | Caspase recruitment domain-containing protein 10 (**Q9BWT7**) | 116.2 | 209 | 74.40 | 87.99 | 41.00 | 64.99 |
| **ZO2** | Tight junction protein ZO-2 (**Q9UDY2**) | 134.2 | 207 | 70.80 | 78.00 | 41.50 | 76.50 |
| **MYO6** | Unconventional myosin-VI (**Q9UM54**) | 150.0 | 206 | 62.40 | 71.00 | 52.00 | 68.00 |
| **IQEC1** | IQ motif and SEC7 domain-containing protein 1 (**Q6DN90**) | 108.5 | 204 | 65.30 | 75.50 | 45.00 | 67.50 |
| **DDB1** | DNA damage-binding protein 1 (**Q16531**) | 127.2 | 202 | 61.30 | 78.00 | 50.00 | 57.00 |
| **CLAP1** | CLIP-associating protein 1 (**Q7Z460**) | 169.8 | 201 | 61.40 | 63.50 | 48.50 | 70.50 |
| **C170B** | Centrosomal protein of 170 kDa protein B (**Q9Y4F5**) | 172.1 | 201 | 67.80 | 74.00 | 36.00 | 73.50 |
| **ZN316** | Zinc finger protein 316 (**A6NFI3**) | 108.7 | 200 | 66.80 | 65.67 | 48.69 | 65.67 |

**Supplementary Table S2.** List of primary and secondary antibodies used in this study

| Peptide/ Protein Target (clone) | Conjugated | Antigen sequence (if known) | Manufacturer and catalog number | Species raised (mono or poly) | Dilution/ Concentration used | Assay |
|---|---|---|---|---|---|---|
| **EEA1 (C45B10)** | | Residues surrounding Ser70 | CST # 3288 | Rb mono | 1:200 | IF |
| **EGFR (D38B1)** | | | CST # 4267 | Rb monol | 1:2000 | WB |
| **EGFR**[a] | | Amino acids 25-645 | R&D # AF231 | Gt poly | 1:1000 | WB |
| **EGFR (15F8)** | | | CST # 4405 | Rb mono | 1:2000 | WB |
| **EGFR PY1068 (1H12)** | | Residues surrounding Tyr1068 | CST # 2236 | Ms mono | 1:2000 | WB |
| **EGFR PY845** | | | SCB # sc-23420 | Rb poly | 1:500 | WB |
| **EGFR PY1148** | | | CST #2237 | Rb poly | 1:2000 | WB |
| **EGFR (R-1)** | AF647 | Amino acids 6-273 | SCB # sc-101 AF647 | Ms mono | 5 – 40 ug/ml | FC |
| **EGFR (D-20)** | | | SCB # sc-31156 | Gt poly | 1:20 | EM |
| **HA (6E2)** | AF488 | YPYDVPD YA | CST # 2350 | Ms mono | 5 – 40 ug/ml | FC |
| **HA (C29F4)** | Magnetic bead | YPYDVPD YA | CST # 11846 | Rb mono | 1:100 | IP |

| | | | | | | |
|---|---|---|---|---|---|---|
| **HA (C29F4)** | Sepharose bead | YPYDVPD YA | CST # 3956 | Rb mono | 1:100 | IP |
| **HA-Fab (3F10)** | FITC | YPYDVPD YA | Roche # 11988506001 | Rt mono | 1:20 | IF |
| **HA-Fab (3F10)** | Biotin | YPYDVPD YA | Roche # 12158167001 | Rt mono | 200 pM | SPT |
| **PY20** | | | SCB # sc-508 | Ms mono | 1:500 | WB |
| **PY20** | HRP | | SCB # sc-508 | Ms mono | 1:500 | Kinase assay |
| **PY99** | | | SCB # sc-7020 | Ms mono | 1:500 | WB |
| **PY99** | HRP | | SCB # sc-7020 | Ms mono | 1:500 | Kinase assay |
| **RON** | | Amino acids 25-956 | R&D # AF691 | Gt poly | 1:100 | IP |
| **RONβ (C-20)[b]** | | | SCB # sc-322 | Rb poly | 1:500 and 1:20 | WB and EM |
| **RONβ (C81H9)** | | Residues surroundin g Lys624 | CST # 2654 | Rb mono | 1:2000 | WB |
| **RONβ (E-3)[c]** | | Amino acids 531-690 | SCB # sc-74588 | Ms mono | 1:500 | WB |
| **RON PY1238/39** | | Residues surroundin g Tyr1238/39 | R&D # AF1947 | Rb poly | 1:2000 | WB |
| **Src (L4A1)** | | Amino acids 1-110 | CST # 2110 | Ms mono | 1:2000 | WB |
| **Src PY416** | | Residues surroundin g Tyr416 | CST # 2101 | Rb poly | 1:2000 | WB |
| **Goat IgG** | IRDye 800CW | | Li-Cor # 926-32214 | Dk | 1:20,0000 | WB |
| **Goat IgG** | 12 nm colloidal gold | | JIR # 705-205-147 | Dk | 1:20 | EM |
| **Mouse IgG** | IRDye 680RD | | Li-Cor # 926-68070 | Gt | 1:20,000 | WB |
| **Mouse IgG** | IRDye 680RD | | Li-Cor # 926-68072 | Dk | 1:20,000 | WB |
| **Rabbit IgG** | IRDye 800CW | | Li-Cor # 926-32211 | Gt | 1:20,000 | WB |
| **Rabbit IgG** | IRDye 680RD | | Li-Cor # 926-68073 | Dk | 1:20,000 | WB |
| **Rabbit IgG-Fab** | AF555 | | TFS # A-21430 | Gt | 1:500 | IF |
| **Rabbit IgG** | 6 nm colloidal gold | | JIR # 711-195-152 | Dk | 1:20 | EM |

CST: Cell Signaling Technologies; R&D: R&D Systems; SCB: Santa Cruz Biotechnology; JIR: Jackson ImmunoResearch; TFS: Thermo Fisher Scientific.
Rb: Rabbit; Gt: Goat; Ms: Mouse; Rt: Rat; Dk: donkey; Gt: Goat; Mono: monoclonal; Poly: polyclonal.
EM: electron microscopy; FC: Flow cytometry; IF: immunofluorescence; IP: immunoprecipitation; SPT: Single particle tracking; and WB: Western Blot.
[a]Used when blotting for EGFR-Δ998.
[b]Discontinued antibody. Remainder of experiments done with CST # 2654.
[c]Used with the PY1238/39 RON antibody.

**Supplementary Table S3.** List of primer and ultramer sequences used in this study.

| | |
|---|---|
| **Ultramer sequence** | CACCATGGAGCTCCTCCCGCCTCAGTCCTTCCTGTTGCTGCTGCTGTTGCCTGA CAAGCCCGCGGCGGGCTATCCTTACGACGTGCCTGACTACGCCGCAGCAGCA GAGGACTGGCAGTGCCCGCACA |
| **RON-K1114M** | Forward: GTGATGCGACTTAGTGACATGATGGCACATTGGATTC<br>Reverse: GAATCCAATGTGCCATCATGTCACTAAGTCGCATCAC |
| **EGFR-KD** | Forward: CGCCGGATCCCCAACGAATGGGCCTAAG<br>Reverse: CGAGGTCGACGGTATCGATAAGCTTTGCTCCAATAAATTCACTGC |
| **EGFR-I682Q** | Forward: CAACCAAGCTCTCTTGAGGCAGTTGAAGGAAACTGAATTC<br>Reverse: GAATTCAGTTTCCTTCAACTGCCTCAAGAGAGCTTGGTTGG |
| **EGFR-V924R** | Forward: GATGTCTACATGATCATGCGCAAGTGCTGGATGATA<br>Reverse: TATCATCCAGCACTTGCGCATGATCATGTAGACATC |
| **EGFR-Δ998** | Forward: GTTAAGCTTGGTACCGAGCTCGGATCCAGTACCCTTCACCATGCGACCCTCCGGGAC<br>Reverse: CCCTCTAGACTCGAGCGGCCGCCTAGAAGTTGGAGTCTGTAGGACTTGGC |

#### SUPPLEMENTARY METHODS

##### Dephosphorylation assay

$HEK^{RON}$ cells were transiently transfected with WT or Δ998-EGFR and allowed to attach and recover overnight. Cells were activated with 50 nM EGF for 2 min followed by 10 μM Afatinib for 20 or 40 sec. Media was removed and reactions were stopped by placing plates on top of a layer of liquid nitrogen. Protein lysates were harvested and quantified by BCA. RON protein was immunoprecipitated from the lysates with anti-RON antibody, and immunoblotted.

##### Mass Spectrometry

$A431^{RON}$ cells were harvested with lysis buffer and immunoprecipitated with an anti-HA antibody. Samples were run in a reducing 4-20% polyacrylamide gel for separation, washed in distilled water for 15 min, and incubated with GelCode Blue stain reagent (Thermo Fisher Scientific, cat # 24590) for 1 hr at RT. Both top and bottom RON bands were excised from the gel and samples sent to the Proteomics Core at UT Southwestern.

### Supplementary Methods

#### Channel Registration

The two color channels used in data collection were registered to one other using a locally weighted mean transform computed from sets of fiducial coordinates collected prior to each experiment. Coordinates of bright spots (control points) in fiducial images from each channel were paired together based on their separation from one another, with pairings greater than 10 pixels ($\approx$ 1.667 micrometers) being thrown out automatically. The selected pairs were then inspected visually to ensure appropriate pairing between the color channels, with pairings deemed inappropriate being thrown out before proceeding. A locally weighted mean transform was then computed from the control points using the MATLAB Image Processing Toolbox [The MathWorks, Inc.] method *fitgeotrans* with the parameter n set to 10. The resulting transform was then applied to coordinates found from raw data in each channel using the MATLAB Image Processing Toolbox [The MathWorks, Inc.] method *transformPointsInverse*.

Estimates of the channel registration errors remaining after the correction procedure were computed as follows. Given a set of control points $(x_{1,i}, y_{1,i}, x_{2,i}, y_{2,i})$ for $i = 1, 2, \ldots, \text{NPairs}$ where the first subscript denotes the channel and the second subscript denotes the pair number, we apply the locally weighted mean corrections to control points from the appropriate channel, e.g., to $(x_{1,i}, y_{1,i})$. The registration error is then defined as the root mean squared error between the corrected control points $(x^*_{1,i}, y^*_{1,i})$ and their associated pair control points $(x_{2,i}, y_{2,i})$ as

$$\sigma_{\text{overlay}} \equiv RMSE = \sqrt{\frac{1}{\text{NPairs}} \sum_{i=1}^{\text{NPairs}} [(x^*_{1,i} - x_{2,i})^2 + (y^*_{1,i} - y_{2,i})^2]}$$

#### Single Particle Tracking

Trajectories are constructed from raw images taken for each color channel using home-built software packages. Emitter coordinates are estimated by fitting the emitter images to a Gaussian point spread function model. Emitter coordinates are stitched together into trajectories using a simplified version of the cost-matrix approach [Jaqaman, 2008] wherein the merge/split process is omitted and the associated costs are specified based on fluorophore blinking kinetics.

#### Hidden Markov Model

Rate parameters governing the transitions between the freely diffusing state (free, non-dimer) and the dimer state were determined based on a two-state hidden Markov model (HMM). Trajectories were classified frame-to-frame as being in either the free state or the dimer state using the Viterbi algorithm using the rate parameters found in the HMM analysis.

### Mathematical Formalism of the HMM Analysis

#### Probability Densities of the Observed Separations

To construct an HMM, we must define the emission density for each model state. The emission density is the probability density of observing a separation given an underlying state of the HMM, e.g., the probability density of observing a separation between two particles given that they were dimerized.

For the free state, we find the desired emission density additionally conditioned on the previous observation, i.e., the density $g(d_n|d_{n-1}, \text{free})$, where $d_n$ is the separation between the two particles in observation $n$ and $d_{n-1}$ is the separation between those same particles in observation $n-1$. We derive the density $g(d_n|d_{n-1}, \text{free})$ by finding the analogous density in Cartesian coordinates $f(\Delta x, \Delta y|\text{free})$ where $\Delta x \equiv \Delta x_n - \Delta x_{n-1}$, $\Delta y \equiv \Delta y_n - \Delta y_{n-1}$, $(\Delta x_n)^2 + (\Delta y_n)^2 = d_n^2$, and $(\Delta x_{n-1})^2 + (\Delta y_{n-1})^2 = d_{n-1}^2$. The quantities $\Delta x$ and $\Delta y$ are both assumed to be Normally distributed with mean zero and variance $\sigma_n^2 = \sigma_{1,n}^2 + \sigma_{2,n}^2 + \sigma_{1,n-1}^2 + \sigma_{2,n-1}^2 + 2(2D)\Delta t_n$ where $\sigma_{1/2,n/n-1}$ is the standard error of the estimates of particle positions 1/2 in observations n/n-1, $D$ is the diffusion constant of each of the particles, and $\Delta t_n \equiv t_n - t_{n-1}$ is the time between observations $n$ and $n-1$. The additional multiplicative factor of 2 in front of the diffusion constant can be understood more readily by deriving $f(\Delta x, \Delta y|\text{free})$ treating one of the particles as fixed, the result being that the moving particle must diffuse with twice its true diffusion constant. The expression $f(\Delta x, \Delta y|\text{free})$ is then converted to polar coordinates $(d_n, \theta)$ yielding $g(d_n, \theta|d_{n-1}, \text{free})$. The desired emission density $g(d_n|d_{n-1}, \text{free})$ is then obtained by integrating $g(d_n, \theta|d_{n-1}, \text{free})$ over $\theta$ assuming that $\theta$ is distributed uniformly on the interval $[0, 2\pi]$, yielding the expression

$$g(d_n|d_{n-1}, \text{free}) = \frac{d_n}{\sigma_n^2} \exp\left(-\frac{d_n^2 + d_{n-1}^2}{\sigma_n^2}\right) I_0\left(\frac{d_n d_{n-1}}{\sigma_n^2}\right)$$

where $I_0(\cdot)$ is the zeroth order modified Bessel function of the first kind. The free state emission density for the first observation is estimated by the expression $g(d_1|d_1, \text{free})$ treating each appearance of $d_1$ as independent observations of the separation in subsequent frames, i.e., setting $\Delta t_n = 1$ frame.

For the dimer state, we proceed with an approach similar to that used to derive the expression for the free state emission density. We begin by finding the probability density $f(\Delta x, \Delta y|\text{dimer})$ where $\Delta x$ and $\Delta y$ are now defined as $\Delta x \equiv \Delta x_n - \Delta x'$ and $\Delta y \equiv \Delta y_n - \Delta y'$, where $\Delta x'^2 + \Delta y'^2 = L_{\text{dimer}}^2$ with $L_{\text{dimer}}$ being the true separation between fluorophores attached to the two constituents of a dimer. The terms $\Delta x$ and $\Delta y$ are treated as Normally distributed random variables with mean 0 and variance $\sigma_n^2 = \sigma_{1,n}^2 + \sigma_{2,n}^2 + \sigma_{\text{overlay}}^2$. The addition of the constant term $\sigma_{\text{overlay}}^2$ in the variance is included as an approximation to the true effect of the channel registration error, which in reality will complicate the derivation of $g(d_n|d_{n-1}, \text{dimer})$ from $g(d_n, \theta|d_{n-1}, \text{dimer})$ by breaking the assumption that $\theta$ is distributed uniformly on the interval $[0, 2\pi]$. In practice, this approximation appears to be of little consequence, as the observed channel registration contribution $\sigma_{\text{overlay}}^2$ is typically an order of magnitude smaller than the sum $\sigma_{1,n}^2 + \sigma_{2,n}^2$. Proceeding following the derivation of the free state emission density, we yield the expression for the dimer state emission density

$$g(d_n|L_{\text{dimer}}, \text{dimer}) = \frac{d_n}{\sigma_n^2} \exp\left(-\frac{d_n^2 + L_{\text{dimer}}^2}{\sigma_n^2}\right) I_0\left(\frac{d_n L_{\text{dimer}}}{\sigma_n^2}\right)$$

#### Estimating Rate Parameters

Rate parameters are estimated by maximizing the likelihood of the observed data with respect to the rate parameters of the HMM. The likelihood of the HMM with a set of rate parameters $\theta$ given the observed data $d = [d_1, d_2, \ldots, d_N]$ takes the following form:

$$L(\theta|d) = \pi \mathbf{T}_1 \mathbf{P}_1 \mathbf{T}_2 \mathbf{P}_2 \ldots \mathbf{T}_N \mathbf{P}_N \cdot [1, 1, 1]^T$$

where $\pi = [g(d_1|L_{\text{dimer}}, \text{dimer}),\ g(d_1|d_1, \text{free})]$, $\mathbf{T}_n$ for $n \in 1, 2, \ldots, N$ is a 2x2 transition matrix whose elements give the transition probabilities for the interstate transitions of the HMM, and $\mathbf{P}_n$ for $n \in 1, 2, \ldots, N$ is a 2x2 diagonal matrix whose elements $p_{11}$ and $p_{22}$ give the emission probability densities of an observation $d_n$ given an underlying dimer state or free state, respectively. The elements $t_{i,j}$ for $i, j \in 1, 2$ of $\mathbf{T}_n$ are given by

$$\begin{aligned}
t_{1,2} &= 1 - \exp\left(\Delta t_n k_{1,2}\right) \\
t_{2,1} &= 1 - \exp\left(\Delta t_n k_{2,1}\right) \\
t_{1,1} &= 1 - t_{1,2} \\
t_{2,2} &= 1 - t_{2,1}
\end{aligned}$$

where $k_{1,2}$ is the transition rate from the dimer state to the free state and $k_{2,1}$ is the transition rate from the free state to the dimer state. The rate parameters $k_{1,2}$ and $k_{2,1}$ are thus found by maximizing the sum of the log-likelihoods over all candidate interactions $\sum_{\text{interactions}} \log L(\theta|d)$. The standard errors of these rate parameter estimates are then estimated from elements $h_{i,i}$ of the Hessian of the negative log-likelihood $\mathrm{H}$ as $\sqrt{h_{i,i}^{-1}}$ where $i = 1$ corresponds to $k_{2,1}$ and $i = 2$ corresponds to $k_{1,2}$.
